# Branched-chain amino acid metabolism is a crucial modulator of cellular senescence

**DOI:** 10.1101/2024.09.10.612139

**Authors:** Yuma Aramaki, Kazuki Irie, Hideru Obinata, Shinya Honda, Takuro Horii, Satoko Arakawa, Aiko Tsuchida, Junki Hoshino, Ryosuke Kobayashi, Takashi Izumi, Izuho Hatada, Shigeomi Shimizu, Yoji A. Minamishima, Akimitsu Konishi

## Abstract

Cellular senescence is a complex stress response that results in the permanent arrest of cell proliferation. The accumulation of senescent cells occurs during aging in living organisms, and contributes to tissue dysfunction. Although there are growing lines of evidence that various metabolic changes occur in senescent cells, the link between cellular metabolism and senescence is not yet fully understood. In this study, we demonstrate that alterations in the metabolism of branched-chain amino acids (BCAAs) play a crucial role in establishing cellular senescence. Furthermore, we identified mitochondrial BCAA transamination as a crucial step in this process. Our findings show that various types of cellular stress lead to a reduction in the expression of BCAA aminotransferase 2 (BCAT2), one of the BCAA catabolic enzymes, resulting in decreased catabolism of BCAAs and reduced synthesis of glutamate. The reduction of BCAA catabolites, together with the consequent limitation in glutathione production from glutamate, triggers cellular senescence. Furthermore, we demonstrate that a reduction in BCAT2 levels alone is sufficient to induce cellular senescence, both in cultured cells and in mice. Additionally, our results demonstrate that aging alters BCAA metabolism in both mice and humans. Our findings provide new insights into the metabolic mechanisms underlying cellular senescence, with a particular focus on the role of BCAAs.

## Introduction

Aging is a process characterized by a reduced ability to maintain homeostasis and regenerative capacity in various tissues. This phenomenon is primarily attributed to the accumulation of senescent cells, which are in a state of irreversible cell cycle arrest induced by diverse types of cellular stress, such as DNA damage, oncogene activation, replicative stress, and telomere shortening ^1,2^. Senescent cells differ from healthy cells not only in their cell cycle status, but also in their metabolic state. For instance, human diploid fibroblasts undergo a metabolic shift towards glycolysis during replicative senescence (RS) ^3–5^. Moreover, oncogene-induced senescence (OIS) and therapy-induced senescence (TIS) have been linked to increased tricarboxylic acid (TCA) cycle activity and mitochondrial oxidative metabolism ^6–8^. Additionally, fatty acid metabolism has been implicated in cellular senescence, with OIS and TIS both enhancing fatty acid oxidation, and consequently increasing oxygen consumption ^9,10^. Furthermore, nicotinamide adenine dinucleotide (NAD) metabolism plays a vital role in senescence, as evidenced by the decreased ratio of oxidized and reduced forms of NAD (NAD^+^/NADH ratio) during RS ^11,12^.

Branched-chain amino acids (BCAAs), i.e., leucine, isoleucine, and valine, are essential amino acids utilized not only as protein building blocks but also as sources of cellular energy. BCAA metabolism plays important roles under both physiological and pathological conditions. Epidemiological studies have found increased circulating BCAA levels in patients with obesity or glucose tolerance disorders ^13–18^. Additionally, dietary BCAA intake has been shown to negatively affect insulin sensitivity in rodent models ^15,19–22^. In contrast to these adverse effects on metabolic health, BCAA supplementation generally improves age-associated disorders. Particularly when started in middle age, BCAA supplementation enhances glucose tolerance, cardiac and skeletal muscle metabolism, and overall survival ^23–25^. However, reports also exist on the detrimental effects of BCAA, such as restricted dietary BCAA intake improving lifespan, whereas a high-BCAA diet reduces lifespan ^26–29^. Consequently, the effect of BCAA on aging remains ambiguous, and the role of BCAA metabolism in cellular senescence remains poorly understood.

In this report, we demonstrate that various types of senescence-inducing stress and aging itself reduce cellular BCAA catabolism, by suppressing the BCAA catabolizing enzyme BCAA aminotransferase 2 (BCAT2), and that BCAT2 plays a crucial role in regulating cellular senescence.

## Results

### Accumulation of intracellular BCAAs in senescent cells

To clarify the metabolic changes that occur in senescent cells, we developed a system to induce cellular senescence by telomere shortening. Telomere shortening is widely recognized as a cause of cellular senescence. TRF2 (telomeric repeat-binding factor 2), a subunit of the shelterin complex, is known to protect telomeres. The N-terminal basic domain of TRF2 prevents resolution of the telomere loop structure, termed the t-loop. The ectopic expression of a TRF2 mutant lacking the basic domain (TRF2ΔB) rapidly induces t-loop-sized deletions in telomeres. Consequently, cells expressing TRF2ΔB enter cellular senescence within a few days ^30,31^ (Supplementary Figure S1A). We established tetracycline-inducible TRF2ΔB (tetTRF2ΔB) cell lines from human telomerase reverse transcriptase (hTERT)-immortalized IMR90 normal human diploid fibroblasts (IMR90-tetTRF2ΔB), together with vector control cells (IMR90-tetOne) (Supplementary Figure S1B). As expected, rapid telomere shortening and persistent DNA damage response were observed at the telomeres of IMR90-tetTRF2ΔB cells following doxycycline (Dox) induction (Supplementary Figure S1C–E). IMR90-tetTRF2ΔB cells demonstrated suppressed proliferation within 3 days, and almost completely halted growth 9 days post-Dox induction (Supplementary Figure S2A, B). Dox-induced growth arrest in IMR90-tetTRF2ΔB cells was confirmed to be cellular senescence using various criteria, including increased senescence-associated β-galactosidase (SA-β Gal) activity, upregulation of the cyclin-dependent kinase inhibitors (CDKNs) p16 and p21, and senescence-associated secretory phenotype (SASP)-related genes, increased formation of senescence-associated heterochromatin foci (SAHF), and loss of lamin B1 (Supplementary Figure S2C–L). This Dox-induced cellular senescence in tetTRF2ΔB cells was highly reproducible and synchronous.

We then used tetTRF2ΔB cells for the metabolomic analysis of intracellular metabolites, using liquid chromatography-tandem mass spectrometry (LC-MS). The intracellular metabolite profiles of tetTRF2ΔB cells showed significant time-dependent changes post-Dox induction (Figure 1A, Supplementary Figure S3A). Metabolite set enrichment analysis demonstrated that various metabolic pathways were altered in senescent cells, when comparing pathways enriched on day 3 post-Dox induction (characterized by telomere shortening, yet incomplete cellular senescence) and on day 9 (characterized by completed telomere shortening and cellular senescence). Whereas there was considerable overlap in most pathways, specific enrichment of BCAA degradation, purine, and butyrate metabolic pathways was noted on day 9 (Figure 1B, C). The BCAA catabolic pathway appeared to be the most relevant as all three BCAAs contributed significantly to the determination of the first and second principal component axes which clearly characterized the metabolomic changes of day 9 in the principal component analysis (Supplementary Fig. S3B, C). We hence shifted our focus to alterations in BCAA metabolism during cellular senescence. Figure 1D demonstrates a gradual increase in all three intracellular BCAAs in IMR90-tetTRF2ΔB cells compared with IMR90-tetOne control cells following Dox induction. Additionally, the accumulation of BCAAs was observed in senescent hTERT-immortalized IMR90 (IMR90-hTERT) cells induced by X-ray irradiation (IR) or chemical agents (Figure 1E, F), as well as in senescent hTERT-immortalized RPE1 normal human retinal pigment epithelial cells (RPE1-hTERT) induced by IR, and in tetracycline-inducible TRF2ΔB cell lines derived from RPE1-hTERT cells (Figure 1G, H, and Supplementary Figure S2M, N). Collectively, these findings suggest that alterations in BCAA metabolism are associated with cellular senescence.

**Figure 1.**
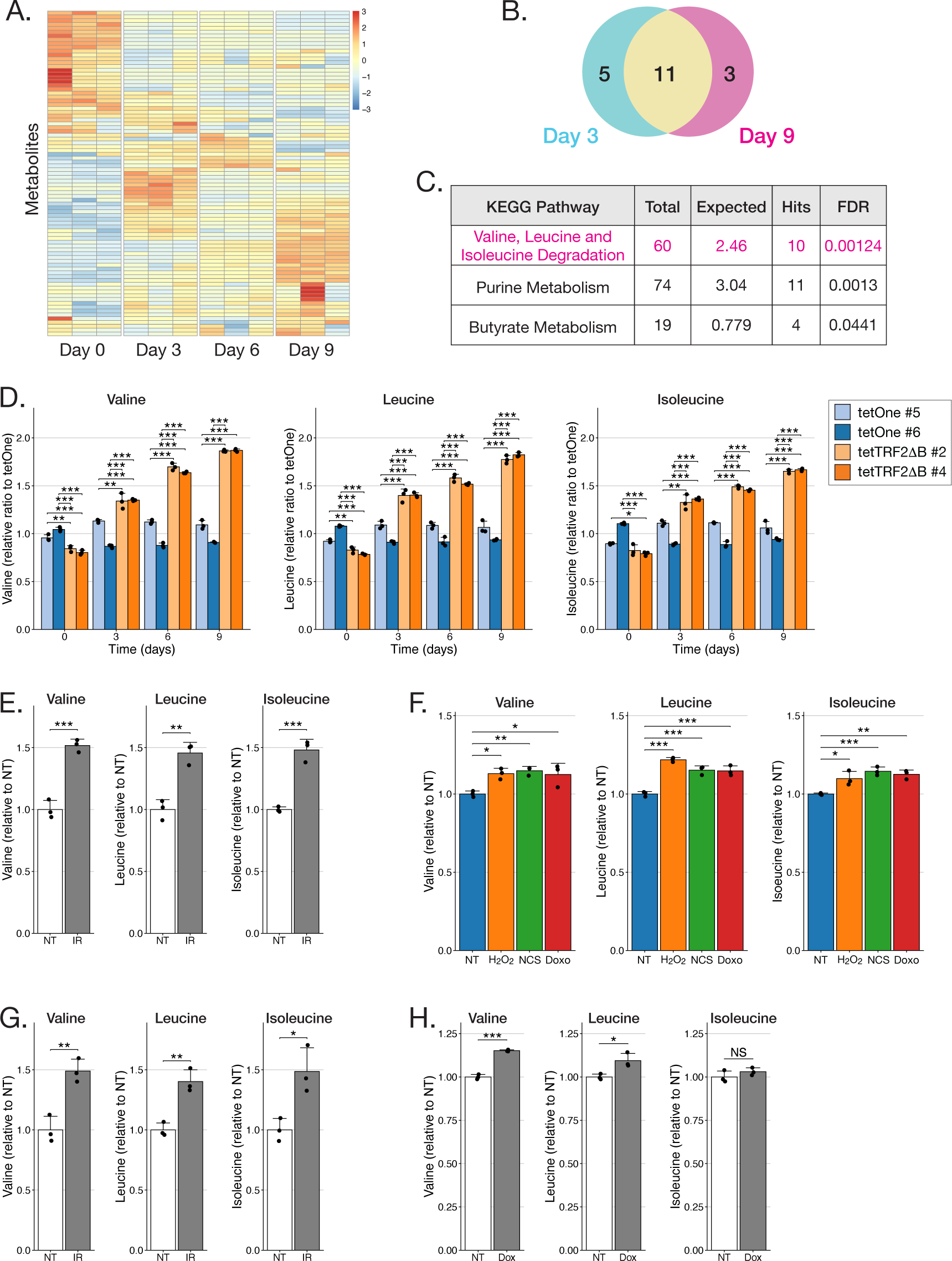
Accumulation of intracellular branched-chain amino acids (BCAAs) in senescent cells. **(A)** Heatmap of intracellular metabolites in doxycycline (Dox)-induced IMR90-tetTRF2ΔB cells. IMR90-tetTRF2ΔB #2 cells were incubated with Dox for the indicated times, and then subjected to metabolome analysis. Data were collected from the biological triplicates at each time point. The heatmap of z-scores is presented. **(B, C)** Metabolite set enrichment analysis of Dox-induced IMR90-tetTRF2ΔB cells. (B) Numbers of enriched Kyoto Encyclopedia of Genes and Genomes (KEGG) metabolic pathways on Day 3 and Day 9 after Dox induction. (C) Detailed list of KEGG pathways enriched only on Day 9. FDR, false discovery rate. **(D)** Accumulation of intracellular BCAAs in Dox-induced IMR90-tetTRF2ΔB cells. IMR90-tetTRF2ΔB or IMR90-tetOne cell clones were incubated with Dox for the indicated times, and then subjected to metabolome analysis. Bars represent the mean relative ratios to control tetOne cells from three independent experiments, and the dots represent the values from each experiment. Error bars indicate the standard deviation (SD). \**p* < 0.05, \*\**p* < 0.01, and \*\*\**p* < 0.001, by two-way ANOVA with Tukey’s test. **(E, F)** Accumulation of intracellular BCAAs in irradiation or chemically induced senescent IMR90-hTERT cells. Cells were treated with 4 Gy X-rays (IR) (E), 800 µM hydrogen peroxide (H_2_O_2_), 200 ng/mL neocarzinostatin (NCS), or 250 nM doxorubicin (Doxo) (F), and incubated for 8 to 14 days to induce cellular senescence, and then subjected to metabolome analysis. Bars represent the mean relative ratios to the cells with no treatment (NT) from three independent experiments, and the dots represent the values from each experiment. Error bars indicate SD. \**p* < 0.05, \*\**p* < 0.01, and \*\*\**p* < 0.001, by the unpaired Student *t*-test (E), and one-way ANOVA with Dunnett’s test (F). **(G)** Accumulation of intracellular BCAAs in X-ray irradiation-induced senescent RPE1-hTERT cells. Cells were irradiated with 4 Gy X-rays (IR), and incubated for 9 days to induce cellular senescence, and then subjected to metabolome analysis. Bars represent the mean relative ratios to NT from three independent experiments, and the dots represent the values from each experiment. Error bars indicate SD. \**p* < 0.05 and \*\**p* < 0.01, by the unpaired Student *t*-test. **(H)** Accumulation of intracellular BCAAs in Dox-treated RPE1-tetTRF2ΔB cells. RPE1-tetTRF2ΔB cells were incubated with or without Dox for 9 days, and then subjected to metabolome analysis. Bars represent the mean relative ratios to NT from three independent experiments, and the dots represent the values from each experiment. Error bars indicate SD. \**p* < 0.05 and \*\*\**p* < 0.001, by the unpaired Student *t*-test. NS, not significant.

### Suppression of BCAA transamination in senescent cells

In mammals, the initial two steps of BCAA metabolism are identical for all three BCAAs. BCAA aminotransferase (BCAT) catalyzes the catabolism of BCAA to synthesize branched-chain α-ketoacids (BCKAs); specifically, α-ketoisocaproic acid (α-KIC), α-ketomethylvaleric acid (α-KMV), and α-ketoisovaleric acid (α-KIV). There are two isozymes of BCAT in mammals: BCAT1, which is localized in the cytoplasm, and BCAT2, a mitochondria-localized enzyme. Subsequently, the BCKAs undergo oxidative decarboxylation by the branched-chain α-ketoacid dehydrogenase (BCKDH) complex, within the mitochondrial matrix (Figure 2A) ^32^.

**Figure 2.**
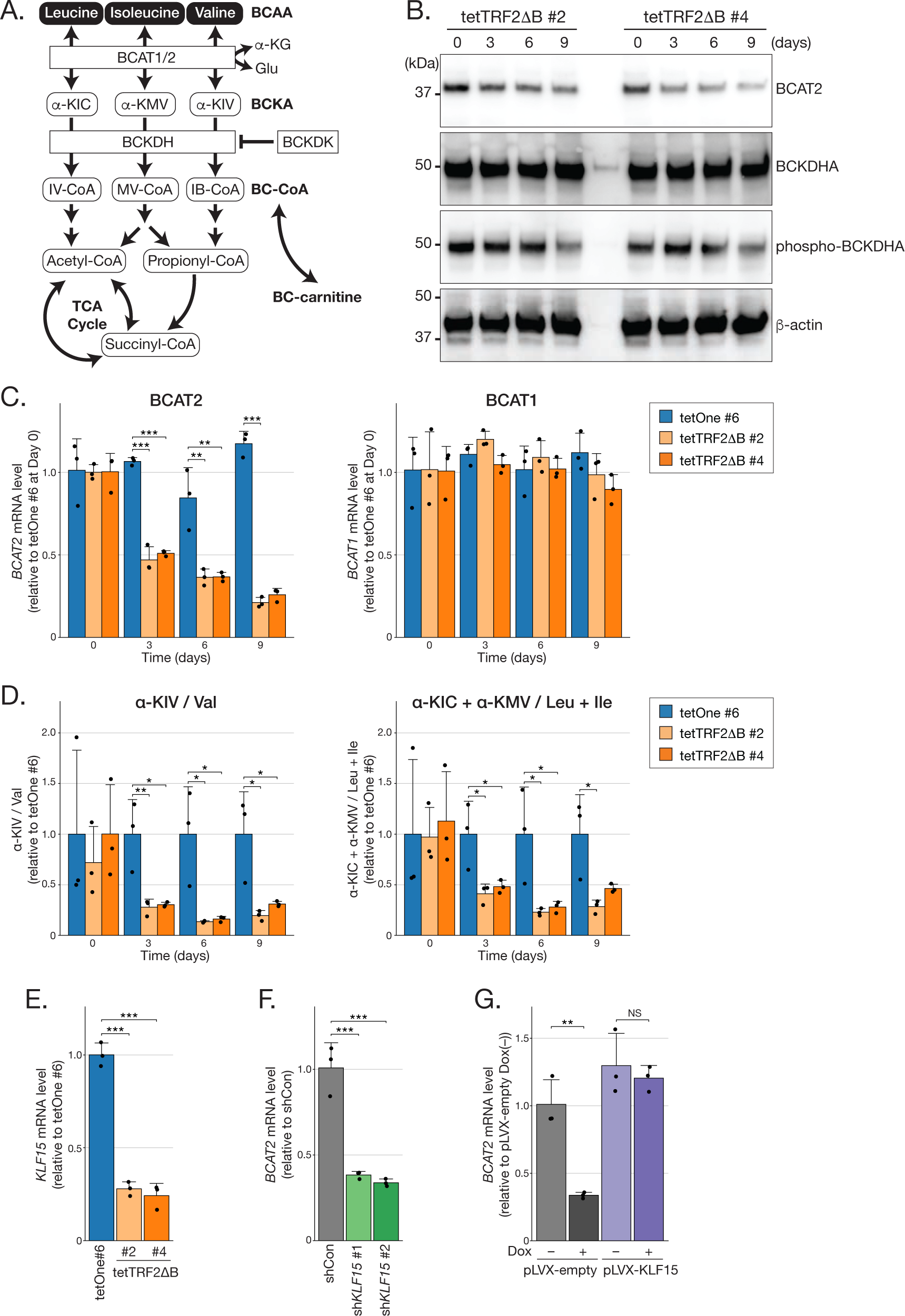
Suppression of BCAA transamination in senescent cells. **(A)** Schematic of BCAA metabolism. BCAT, BCAA aminotransferase; BCKDH, branched-chain α-ketoacid dehydrogenase; BCKDK, BCKDH kinase; α-KG, α-ketoglutarate; α-KIC, α-ketoisocaproic acid; α-KMV, α-ketomethylvaleric acid; α-KIV, α-ketoisovaleric acid; IV-CoA, isovaleryl-coenzyme A (CoA); MV-CoA, α-methylbutanoyl-CoA; IB-CoA, isobutyryl-CoA; BCKA, branched-chain α-ketoacid; BC-CoA, branched-chain acyl-CoA; BC-carnitine, branched-chain acyl-carnitine. **(B, C)** Suppression of BCAT2 expression in Dox-induced tetTRF2ΔB cells. IMR90-tetTRF2ΔB or IMR90-tetOne cell clones were incubated with Dox for the indicated times, and then subjected to immunoblotting (B) or RT-qPCR (C). β-Actin was used as a loading control (B). Data are normalized to β-actin, and the relative ratios to IMR90-tetOne clone #6 without Dox are presented (C). Bars represent the mean values from three independent experiments, and the dots represent the values from each experiment. Error bars indicate SD. \*\**p* < 0.01 and \*\*\**p* < 0.001, by two-way ANOVA with Dunnett’s test. **(D)** Decreased BCKA to BCAA ratio in Dox-treated tetTRF2ΔB cells. IMR90-tetTRF2ΔB and IMR90-tetOne cell clones were incubated with Dox for the indicated times, and then subjected to metabolome analysis. Ratios of keto-valine to valine (α-KIV/Val), and the combined ratio of keto-leucine and keto-isoleucine to the sum of leucine and isoleucine (α-KIC+α-KMV/Leu+Ile) are presented. Bars represent the mean relative ratios to IMR90-tetOne clone #6 from three independent experiments, and the dots represent the values from each experiment. Error bars indicate SD. \*\**p* < 0.01 and \*\*\**p* < 0.001 by two-way ANOVA with Dunnett’s test. **(E)** Reduced KLF15 expression in Dox-treated tetTRF2ΔB cells. IMR90-tetTRF2ΔB and IMR90-tetOne cell clones were incubated with Dox for 9 days, and then subjected to RT-qPCR. Data are normalized to β-actin, and the ratios relative to tetOne clone #6 are presented. Bars represent the mean values from three independent experiments, and the dots represent the values from each experiment. Error bars indicate SD. \*\*\**p* < 0.001, by one-way ANOVA with Dunnett’s test. **(F, G)** Regulation of BCAT2 expression by KLF15. (F) IMR90-hTERT cells were lentivirally introduced with two independent shRNAs for *KLF15* (sh*KLF15* #1, #2) and an shRNA control (shCon), and then subjected to RT-qPCR to detect BCAT2. (G) IMR90-tetTRF2ΔB cells infected lentivirally with the KLF15 expression vector (pLVX-KLF15) or empty vector (pLVX-empty) were incubated with Dox for 9 days, and then subjected to RT-qPCR to detect BCAT2. Data are normalized to β-actin, and the ratios relative to shCon (F) and to pLVX-empty without Dox (G) are presented. Bars represent the mean values from three independent experiments, and the dots represent the values from each experiment. Error bars indicate SD. \*\**p* < 0.01 and \*\*\**p* < 0.001, by one-way ANOVA with Dunnett’s test (F) and two-way ANOVA with Tukey’s test (G). NS, not significant.

Observation of the accumulation of all three BCAAs in senescent cells led us to hypothesize that alterations in BCAT or BCKDH have occurred in these cells. Consistent with our hypothesis, we observed a marked decrease in BCAT2 expression at both the protein and mRNA levels during cellular senescence in Dox-treated tetTRF2ΔB cells. In contrast, the expression of BCAT1 and BCKDH remained unaffected (Figure 2B, C). BCKDH activity is primarily regulated by post-translational modification, in which BCKDH kinase (BCKDK) phosphorylates the E1α subunit of BCKDH (BCKDHA) to inhibit its activity (Figure 2A). We found that although total BCKDHA protein levels remained constant, there was a decrease in BCKDHA phosphorylation during senescence, suggesting an upregulation of BCKDH activity (Figure 2B). Further supporting these findings, the ratio of keto-valine to valine (α-KIV/Val), and the combined ratio of keto-leucine and keto-isoleucine to the sum of leucine and isoleucine (α-KIC+α-KMV/Leu+Ile) were decreased in Dox-treated IMR90-tetTRF2ΔB cells compared with control cells (Figure 2D). It was technically difficult to distinguish between α-KIC and α-KMV by LC-MS analysis. Therefore, these compounds were evaluated in combination. Taken together, these results indicate that the downregulation of BCAA catabolism in senescent cells likely occurs at the transamination step, and may be attributed to the decreased activity of BCAT2.

Several factors are known to regulate *BCAT2* transcription, including peroxisome proliferator-activated receptor γ coactivator 1α (*PGC-1α*), Krüppel-like factor 15 (*KLF15*), and sterol regulatory element-binding transcription factor 1 (*SREBF1*) ^33–35^. Our analysis of these factors demonstrated a reduction in *KLF15* expression, but not *PGC-1α* or *SREBF1*, in Dox-treated IMR90-tetTRF2ΔB cells (Figure 2E, and Supplementary Figure S4A, B). The knockdown of KLF15 inhibited *BCAT2* mRNA expression, and the exogenous expression of KLF15 was able to rescue the transcriptional suppression of *BCAT2* in Dox-treated IMR90-tetTRF2ΔB cells (Figure 2F, G and Supplementary Figure S4C, D). These results suggest that KLF15 plays a regulatory role in BCAT2 expression during cellular senescence.

### Regulation of cellular senescence by BCAT2

As BCAT2 expression was found to be reduced in senescent cells, we next investigated whether a decrease in BCAT2 expression could induce cellular senescence. BCAT has two isozymes, BCAT1 and BCAT2, but the knockdown of only BCAT2 by short hairpin RNA (shRNA) increased all three intracellular BCAAs and decreased the ratio of BCKA to BCAA in IMR90-hTERT cells, similar to the effects observed upon the induction of senescence (Figure 3A, Supplementary Figure S5A, B). BCAT2 knockdown nearly completely inhibited cell proliferation (Figure 3B). To rule out off-target effects of shRNA, we conducted a rescue experiment using an shRNA-resistant BCAT2 expression vector, which completely reversed the inhibition of proliferation (Figure 3B). Concurrently, BCAT2 knockdown increased SA-β Gal activity, upregulated p16, p21, and SASP-related gene expression, and decreased lamin B1 expression, indicating that a reduction in BCAT2 expression on its own can induce cellular senescence (Figure 3C–H). BCAT2 knockdown similarly induced senescence in RPE1-hTERT cells (Supplementary Figure S5C). Furthermore, the BCAT inhibitor gabapentin was found to inhibit cell proliferation in IMR90-hTERT cells (Supplementary Figure S5D).

**Figure 3.**
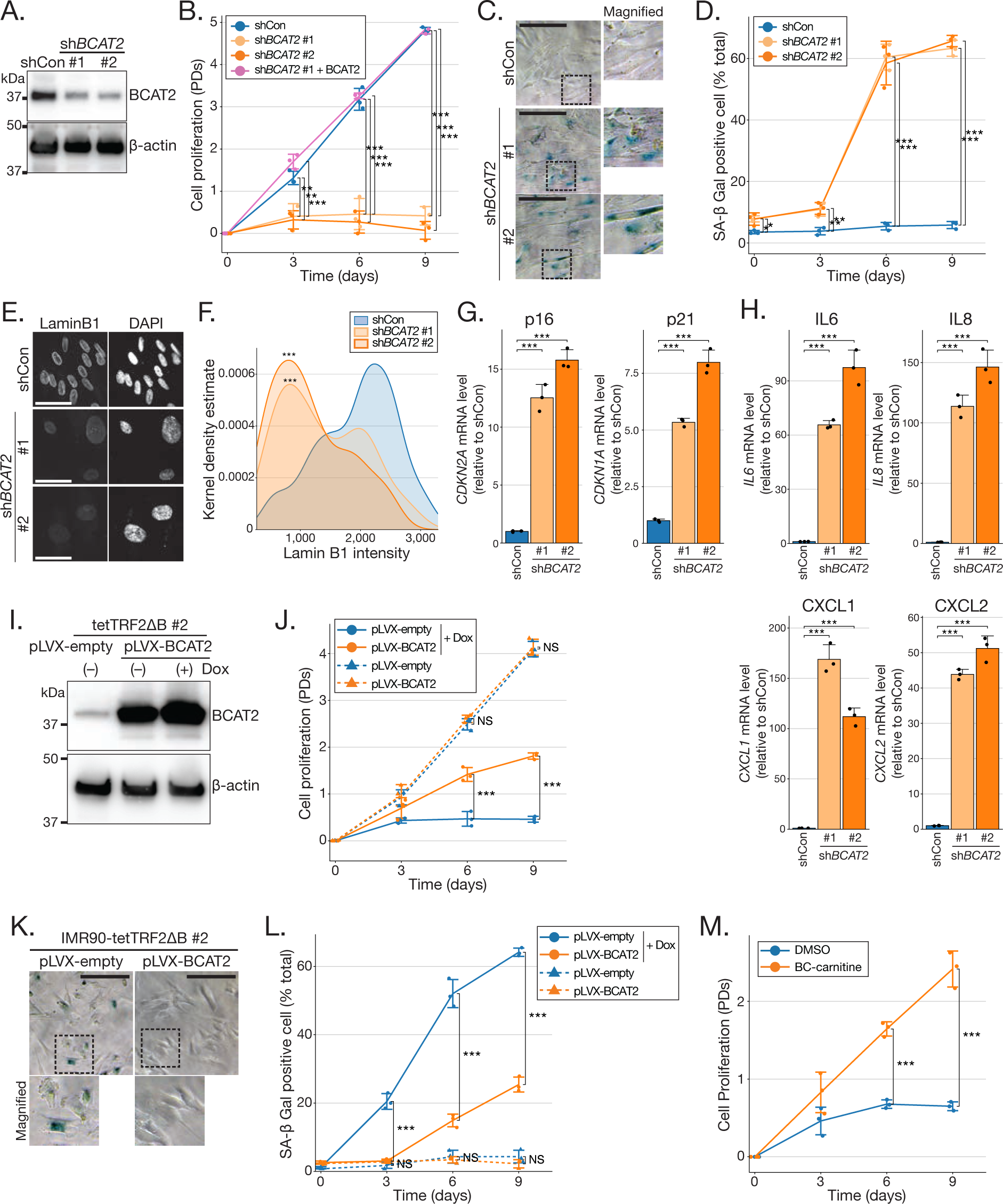
Regulation of cellular senescence by BCAT2. **(A)** Immunoblot analysis of BCAT2 knockdown efficiency. IMR90-hTERT cells were lentivirally introduced with either of two independent shRNAs for *BCAT2* (sh*BCAT2* #1, #2) or the shRNA control (shCon), and then subjected to immunoblotting to detect the BCAT2 protein. β-actin was used as a loading control. **(B)** Inhibition of cell proliferation by BCAT2 knockdown. IMR90-hTERT cells lentivirally introduced with shCon or sh*BCAT2* were incubated for the indicated times, and cumulative population doublings (PDs) were calculated. Complementation of BCAT2 was performed by lentiviral infection of BCAT2 cDNA expression vector (pLVX-BCAT2) before the introduction of sh*BCAT2* #1 targeting the 3′-UTR (untranslated region) of *BCAT2* mRNA (sh*BCAT2* #1 + BCAT2). Data are presented as mean values from three independent experiments, and the dots represent the values from each experiment. Error bars indicate SD. \*\**p* < 0.01 and \*\*\**p* < 0.001, by two-way ANOVA with Tukey’s test. **(C, D)** Increase of SA-β Gal activity by BCAT2 knockdown. (C) Representative images of SA-β Gal staining of *BCAT2*-knockdown cells. IMR90-hTERT cells lentivirally introduced with shCon or sh*BCAT2* were incubated for 9 days after puromycin selection for shRNA-introduced cells, and then subjected to the SA-β Gal staining assay. Images on the right are magnified images of the boxed regions. Scale bars, 100 µm. (D) Quantification of the SA-β Gal-positive cells from the experiment shown in (C). Data are presented as mean values from three independent experiments, and the dots represent the values from each experiment. Error bars indicate SD. \**p* < 0.05, \*\**p* < 0.01, and \*\*\**p* < 0.001, by two-way ANOVA with Dunnett’s test. **(E, F)** Loss of lamin B1 in *BCAT2*-knockdown cells. (E) Representative images of lamin B1 immunofluorescence (IF) of *BCAT2*-knockdown cells. IMR90-hTERT cells were incubated for 9 days after shRNA introduction, and then subjected to IF to detect lamin B1. Scale bars, 50 µm. (F) Quantification of lamin B1 fluorescence intensity at the nuclear periphery, from the experiment shown in (E). \*\*\**p* < 0.001; versus shCon, by one-way ANOVA with Dunnett’s test. **(G, H)** Increased expression of *CDKN*s and SASP-related genes in *BCAT2*-knockdown cells. Total RNAs were isolated from IMR90-hTERT cells 9 days after shRNA introduction, and analyzed by RT-qPCR to detect *CDKN2A* (*p16*), *CDKN1A* (*p21*), *IL6*, *IL8*, *CXCL1*, and *CXCL2* mRNA. Data are normalized to β-actin, and the relative ratios to shCon are presented. Bars represent the mean values from three independent experiments, and the dots represent the values from each experiment. Error bars indicate SD. \*\*\**p* < 0.001, by one-way ANOVA with Dunnett’s test. **(I–L)** Inhibition of cellular senescence by BCAT2 overexpression. (I) Immunoblot analysis of BCAT2 overexpression. IMR90-tetTRF2ΔB #2 cells were infected lentivirally with pLVX-BCAT2 or empty vector (pLVX-empty), and then subjected to immunoblotting to detect the BCAT2 protein. β-Actin was used as a loading control. (J) Recovery of cell growth arrest by BCAT2 overexpression in Dox-treated tetTRF2ΔB cells. IMR90-tetTRF2ΔB #2 cells infected lentivirally with pLVX-BCAT2 or pLVX-empty were incubated with Dox for the indicated times, and cumulative PDs were calculated. Data are presented as mean values from three independent experiments, and the dots represent the values from each experiment. Error bars indicate SD. \*\*\**p* < 0.001, by two-way ANOVA with Tukey’s test. NS, not significant. (K) Representative images of SA-β Gal staining of BCAT2-overexpressing Dox-treated tetTRF2ΔB cells. IMR90-tetTRF2ΔB #2 cells infected lentivirally with pLVX-BCAT2 or pLVX-empty were incubated with Dox for 9 days, and then subjected to the SA-β Gal staining assay. Images at the bottom are magnified images of the boxed regions. Scale bars, 100 µm. (L) Quantification of SA-β Gal-positive cells in the experiment shown in (K). Data are presented as mean values from three independent experiments, and the dots represent the values from each experiment. Error bars indicate SD. \*\*\**p* < 0.001, by two-way ANOVA with Tukey’s test. NS, not significant. **(M)** Abrogation of growth arrest by BC-carnitine supplementation in Dox-treated tetTRF2ΔB cells. IMR90-tetTRF2ΔB #2 cells were induced by Dox in a medium containing 30 µM BC-carnitines for the indicated times, and cumulative PDs were calculated. Data are presented as mean values from three independent experiments, and the dots represent the values from each experiment. Error bars indicate SD. \*\*\**p* < 0.001, by two-way ANOVA with Dunnett’s test.

We next investigated whether the overexpression of BCAT2 could prevent cellular senescence. Indeed, exogenous BCAT2 overexpression suppressed cellular senescence in Dox-treated tetTRF2ΔB cells (Figure 3I–L). Both knockdown and overexpression experiments indicated that BCAT2 activity is crucial for maintaining cell proliferation and preventing cellular senescence. There are two possible explanations as to how BCAT2 regulates cellular senescence. One possibility is that the accumulated intracellular BCAAs exert an inhibitory effect on cell proliferation, leading to senescence. The other is that the shortage of BCAA catabolites, such as BCKAs, induces cellular senescence. To distinguish between these possibilities, we first evaluated the effect of excess BCAAs on cell proliferation. Our results showed that high levels of BCAAs did not affect cell growth, suggesting that the first hypothesis is less likely (Supplementary Figure S5E).

Conversely, an excess amount of BCAAs partially rescued cell growth in Dox-treated IMR90-tetTRF2ΔB cells (Supplementary Figure S5F), indicating that the improved metabolic flux of BCAAs, leading to increased BCKAs or other downstream metabolites, might delay cellular senescence. We hence next hypothesized that BCKAs or downstream metabolites could more effectively suppress senescence. To test this hypothesis, we used carnitine derivatives of BCKA for efficient mitochondrial delivery. Branched-chain acyl-carnitines (BC-carnitines), such as isobutyryl-carnitine, isovaleryl-carnitine, and methylbutyryl-carnitine, can be incorporated into mitochondria and interconverted with downstream BCKA metabolites ^36–38^ (Figure 2A). BC-carnitines abrogated the growth arrest in Dox-treated IMR90-tetTRF2ΔB cells more effectively than BCAAs (Figure 3M). These findings indicate that reduced BCAT2 expression and the subsequent shortage of BCAA catabolites contribute substantially to cellular senescence.

### Induction of cellular senescence by decreased glutathione levels in BCAT2-knockdown cells

Although BC-carnitine alleviated growth arrest in *BCAT2*-knockdown cells, it was unable to reverse it completely (Figure 4E). This implies that a shortage of BCAA catabolites is not the only factor involved in BCAT2-regulated cellular senescence. Because BCAT2 transamination normally uses α-ketoglutarate (α-KG) as the nitrogen acceptor (Figure 2A) ^32^, decreased BCAT2 activity may limit glutamate production from α-KG. Since glutamate is a primary precursor of glutathione (GSH), an essential intracellular antioxidant, decreased BCAT2 activity might exacerbate oxidative stress, which is a primary cause of cellular senescence. In support of this hypothesis, BCAT2 knockdown decreased intracellular glutamate and glutathione levels in IMR90-hTERT cells, but the ratio of reduced GSH to oxidized GSH (GSSG/GSH) was unchanged (Figure 4A, B). These results predicted a decrease in intracellular antioxidant activity. As predicted, BCAT2 knockdown increased intracellular reactive oxygen species (ROS) levels (Figure 4C, D). Supplementation of GSH reduced the intracellular accumulation of ROS, and reversed growth arrest in *BCAT2*-knockdown cells (Figure 4C–E). In contrast, BC-carnitine suppressed cellular senescence in *BCAT2*-knockdown cells, without significantly affecting ROS levels. Combining GSH and BC-carnitine supplementation almost completely abrogated cell growth arrest induced by BCAT2 knockdown, suggesting that glutathione and BCAA catabolites suppress BCAT2 knockdown-induced senescence by different mechanisms (Figure 4E). N-acetyl-L-cysteine (NAC) did not inhibit cellular senescence in *BCAT2*-knockdown cells, possibly because intracellular cysteine and glycine levels did not decrease (Supplementary Figure S6A–C). These results suggest that BCAT2 suppresses cellular senescence through two independent mechanisms, i.e., maintaining BCAA catabolism and maintaining intracellular glutathione levels via glutamate production (Figure 6F).

**Figure 4.**
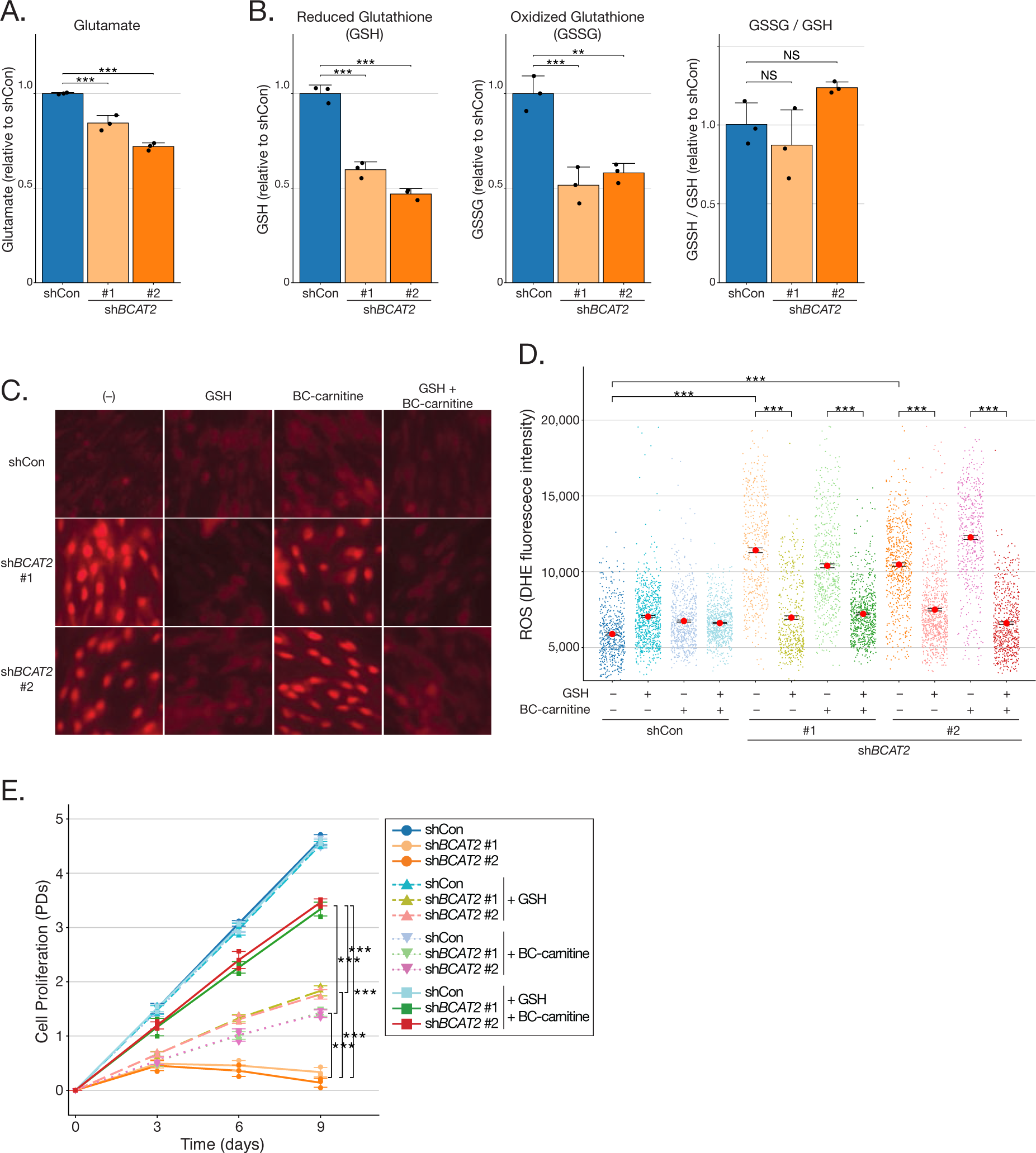
Induction of cellular senescence by decreased glutathione in *BCAT2*-knockdown cells. **(A,B)** Decreased intracellular glutamate and glutathione levels in *BCAT2*-knockdown cells. Metabolites were extracted from IMR90-hTERT cells 3 days after lentiviral shRNA introduction, and then subjected to metabolome analysis. Bars represent the mean relative ratios to shCon from three independent experiments, and the dots represent the values from each experiment. Error bars indicate SD. \*\*\**p* < 0.001, by one-way ANOVA with Dunnett’s test. NS, not significant. **(C, D)** Increased intracellular ROS in *BCAT2*-knockdown cells. (C) Representative images of intracellular ROS. IMR90-hTERT cells lentivirally introduced with shCon or shBCAT2 were incubated for 3 days with 5 mM GSH, 30 µM BC-carnitines, or both, and then subjected to the intracellular ROS detection assay using dihydroethidium (DHE). Oxidized DHE fluorescence images are presented. (D) Quantification of intracellular ROS in the experiment in (C). Red dots indicate mean values. Error bars indicate the standard error (SE). \*\*\**p* < 0.001, by two-way ANOVA with Tukey’s test. **(E)** Abrogation of growth arrest by BC-carnitine supplementation and GSH in *BCAT2*-knockdown cells. IMR90-hTERT cells introduced with shCon or shBCAT2 were incubated with 5 mM GSH, 30 µM BC-carnitines, or both for the indicated times, and cumulative PDs were calculated. Data are presented as mean values from three independent experiments, and the dots represent the values from each experiment. Error bars indicate SD. \*\*\**p* < 0.001, by two-way ANOVA with Tukey’s test.

### Accumulation of senescent cells in Bcat2-deficient mice

To determine the physiological consequences of disrupted BCAA transamination *in vivo*, we generated a full-body *Bcat2*-knockout (KO) mouse using the clustered regularly interspaced short palindromic repeat (CRISPR)-Cas9 system (Supplementary Figure S7A–C). We found that homozygous *Bcat2*-KO mice have low body weight, increased mortality, and severe atrophy in their skeletal muscle and white adipose tissue, consistent with previous reports (Figure 5A, H, Supplemental Figure S7D, E) ^39,40^. No major histological abnormalities other than atrophy were observed in the skeletal muscle and adipose tissue (Figure 5B, C, I, L). As the lower limb reflexes of *Bcat2*-KO mice were normal, it was unlikely that the cause of the muscle atrophy was neurological (Supplemental Figure S7F). We hence hypothesized that the tissue atrophy might be a result of cellular senescence. The decreased expression of lamin B1 and increased expression of p21 and SASP-related genes in the skeletal muscle of *Bcat2*-KO mice supported this hypothesis (Figure 5D–G, Supplemental Figure S8A). There was also a strong increase in SA-β Gal activity in the adipose tissue of *Bcat2*-KO mice (Figure 5I–K, M, Supplemental Figure S8D). Whereas the upregulation of CDKNs and cytokines was unclear, other SASP factors, such as matrix metalloproteinases (MMPs), were significantly increased in the adipose tissue of *Bcat2*-KO mice (Supplemental Figure S8B, C). Although the interstitial spaces of the adipose tissue were increased in *Bcat2*-KO mice, almost all SA-β Gal-positive cells were adipocytes, and not infiltrating inflammatory cells (Figure 5I, Supplemental Figure S8D). Furthermore, most cells with decreased lamin B1 expression were myocytes (Figure 5D). These results indicate that the disruption of BCAT2 contributes to the induction of cellular senescence *in vivo*.

**Figure 5.**
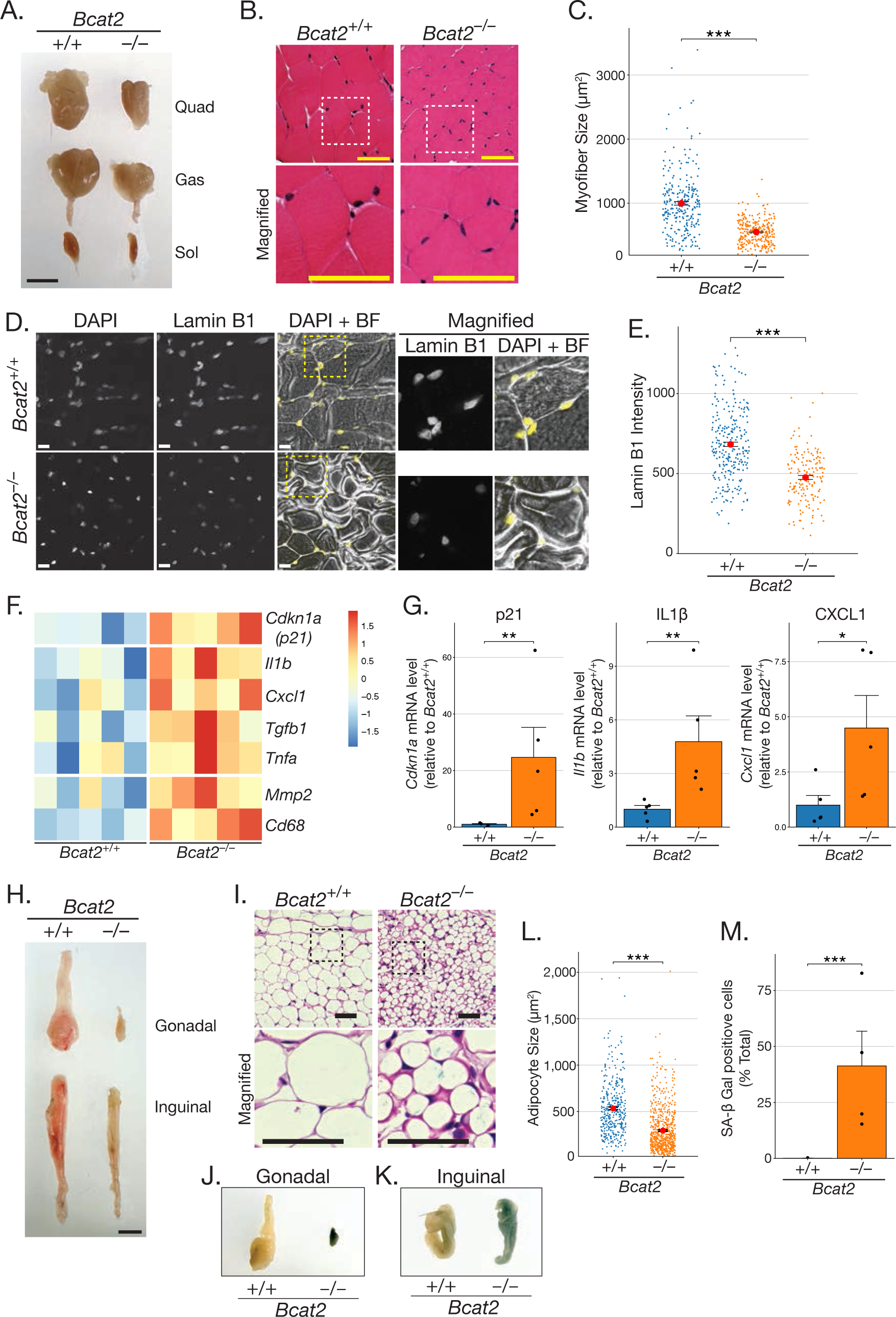
Accumulation of senescent cells and tissue atrophy in *Bcat2*-deficient mice. **(A–C)** Skeletal muscle atrophy in *Bcat2*-knockout (KO) mice. (A) Representative images showing skeletal muscles from age-matched control and *Bcat2*-KO mice (13 weeks old). Quad: quadriceps muscle, Gas: gastrocnemius muscle, Sol: soleus muscle. Scale bar, 5 mm. (B) Representative hematoxylin and eosin (H&E) staining images of the gastrocnemius muscle from age-matched control and *Bcat2*-KO mice (11 weeks old). Images at the bottom are magnified images of the boxed regions. Scale bar, 50 µm. (C) Quantification of myofiber size of the gastrocnemius muscle. Red dots indicate mean values. Error bars indicate SE. \*\*\**p* < 0.001, by the unpaired Student *t*-test. **(D, E)** Decreased lamin B1 expression in the myocytes of *Bcat2*-KO mice. (D) Representative images of lamin B1 immunofluorescence in the gastrocnemius muscle of age-matched control and *Bcat2*-KO mice (11 weeks old). Magnified images of the boxed areas are also presented. Scale bar, 20 µm. BF, bright field. (E) Quantification of lamin B1 intensity in the experiment shown in (D). Red dots indicate mean values. Error bars indicate SE. \*\*\**p* < 0.001, by the unpaired Student *t*-test. **(F, G)** Increased expression of *CDKN* and SASP-related genes in skeletal muscle of *Bcat2*-KO mice. Total RNAs were isolated from the gastrocnemius muscle of age-matched control and *Bcat2*-KO mice (10–13 weeks old). Data are normalized to β-actin, and the heatmaps of z-scores (F) and the relative ratio to *Bcat2*^+/+^ mice (G) are presented. Data are presented as mean values, and the dots represent the values from each mouse (n = 5). Error bars indicate SE. \**p* < 0.05, \*\**p* < 0.01, comparing between *Bcat2*^+/+^ and *Bcat2*^−/−^ using the Wilcoxon rank sum test. **(H–M)** Atrophy and increased SA-β Gal activity in adipose tissues of *Bcat2*-KO mice. (H) Representative images show adipose tissues from age-matched control and *Bcat2*-KO mice (11 weeks old). Gonadal: gonadal adipose tissue, Inguinal: inguinal subcutaneous adipose tissue. Scale bar, 5 mm. (I) Representative histological images of SA-β Gal (blue staining) and H&E staining of gonadal adipose tissues from age-matched control and *Bcat2*-KO mice (11 weeks old). The bottom images are magnified images of the boxed areas. Scale bar, 50 µm. (J, K) Representative macroscopic images of SA-β Gal staining of gonadal adipose tissue (J) and inguinal subcutaneous adipose tissue (K) from age-matched control and *Bcat2*-KO mice (11 weeks old). (L) Quantification of adipocyte size of the gonadal adipose tissues. Red dots indicate mean values. Error bars indicate SE. \*\*\**p* < 0.001, by the unpaired Student *t*-test. (M) Quantification of SA-β Gal positive adipocytes in the experiment shown in (I). Bars represent the mean values, and the dots represent the values from each mouse (n = 4, 10–11 weeks old). Error bars indicate SE. \*\*\**p* < 0.001, by the unpaired Student *t*-test.

### Age-dependent decrease in BCAA catabolism

Because cellular stress, such as telomere shortening, suppressed BCAT2 expression and BCAA catabolism in cultured cells, we hypothesized that aging might affect BCAA catabolism *in vivo*. Indeed, the levels of all three BCAAs increased with age in mouse plasma (Figure 6A). The expression of *Bcat2* mRNA significantly decreased with age in gonadal and inguinal subcutaneous adipose tissues in mice (Figure 6B–D). The production of BCKA in mouse adipose tissue also decreased with age (Figure 6E). Moreover, in a human cohort, the plasma concentrations of all three BCKAs decreased with age (Supplemental Figure S9). These findings indicate that aging reduces the catabolic activity of BCAAs at the transamination step *in vivo* (Figure 6F).

**Figure 6.**
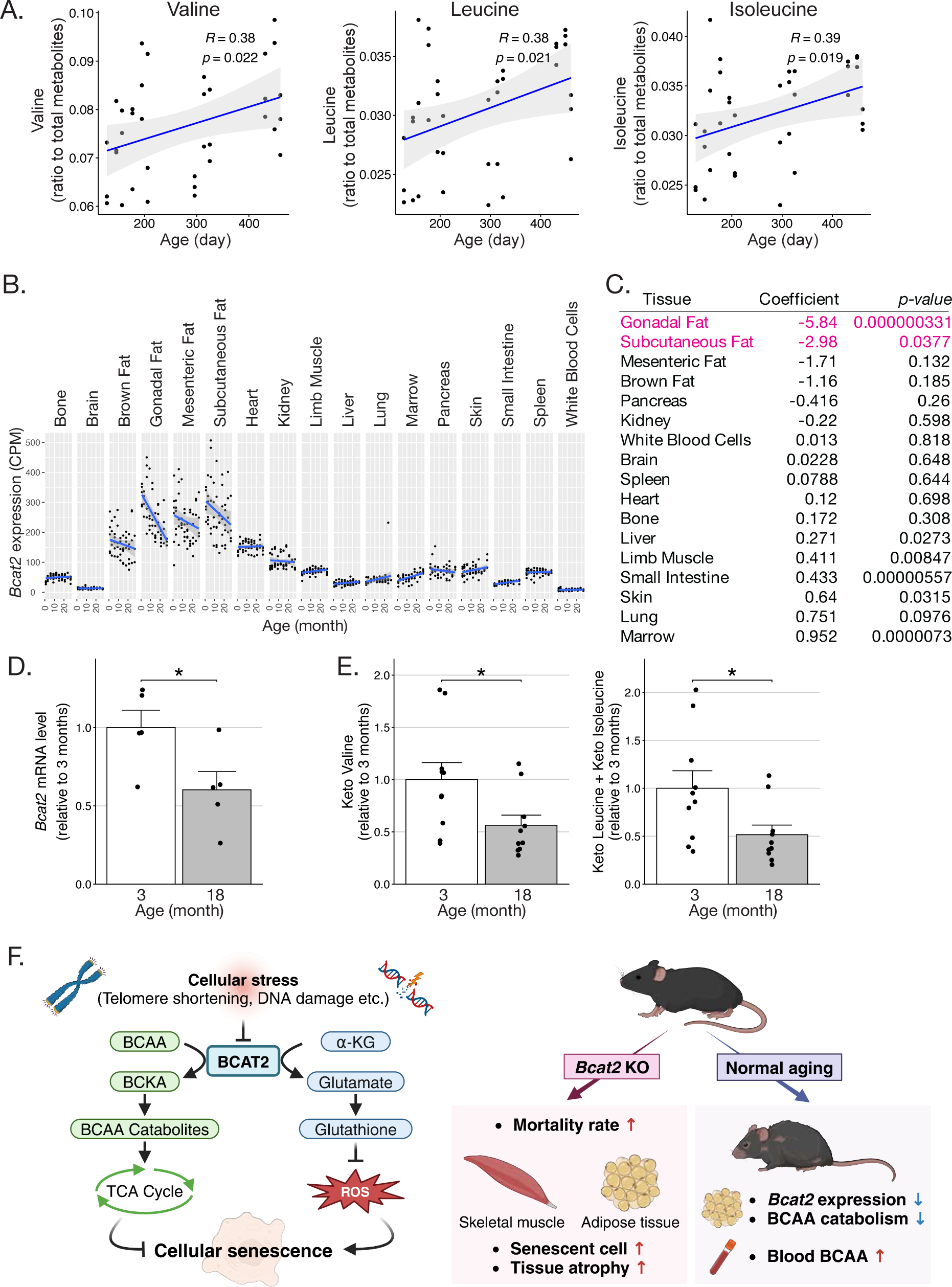
Age-dependent alterations of BCAA metabolism. **(A)** Linear regression analysis between plasma BCAAs and mouse age. Pearson’s correlation coefficients and *p*-values are presented at the top of the plots. The blue lines represent regression lines. The shaded areas show the 95% confidence intervals (CIs). **(B, C)** Linear regression analysis of *Bcat2* mRNA expression in mouse tissues. (B) *Bcat2* read counts from the Tabula Muris Senis Bulk dataset were normalized by calculating counts per million (CPM) and subjected to linear regression analysis. The blue lines represent regression lines. The shaded areas show the 95% CIs. (C) Pearson’s coefficients and statistical values of the data in (B). Tissues indicated in magenta have a significantly negative correlation with age. **(D, E)** Deceased *Bcat2* expression and BCKA production in adipose tissue of aged mice. Inguinal subcutaneous adipose tissues were collected from the indicated months-old normal mice. Data were normalized to β-actin (D) or the total amount of all metabolites (E), and the ratios relative to 3 months-old mice are presented. Data are presented as mean values, and the dots represent the values from each mouse (n = 5 (D), n = 10 (E)). Error bars indicate SE. \**p* < 0.05, by the unpaired Student *t*-test. **(F)** Proposed working model of cellular senescence regulation by BCAT2, and graphical summary of *BCAT2*-KO in mice and age-dependent alterations in BCAA metabolism. BCAT2 mitigates cellular senescence through two distinct mechanisms: the catabolism of BCAAs and the maintenance of intracellular glutathione levels via the production of glutamate (left). The disruption of *BCAT2* in mice results in the accumulation of senescent cells and subsequent tissue atrophy, particularly in skeletal muscle and adipose tissue. Additionally, aging downregulates *BCAT2* expression, leading to alterations in BCAA catabolism in adipose tissue (right). The figure was created using BioRender (https://biorender.com).

## Discussion

In this study, we clarified a previously unknown link between BCAA metabolism and cellular senescence. In normal human cells, various types of senescence-inducing stress increase the intracellular concentration of BCAAs. Among the BCAA-catabolizing enzymes, mitochondrial BCAT2 is downregulated in senescent cells under the control of KLF15. For several reasons, we posit that mitochondrial BCAT2 is a pivotal enzyme for modulating cellular senescence. First, among the metabolic enzymes common to all three BCAAs, BCAT2 was the only enzyme downregulated in senescent cells. Second, although BCAT2 knockdown on its own induced cellular senescence, its overexpression suppressed it. Third, the catabolites of BCAAs downstream of BCAT2 had an inhibitory effect on cellular senescence. Last, the disruption of *BCAT2* in mice resulted in cellular senescence and severe atrophy in their skeletal muscle and adipose tissue.

The mechanisms by which BCAT2 regulates cellular senescence are unique. BCAT2 catalyzes BCAA transamination and glutamate synthesis. BCAT2 knockdown decreases intracellular glutathione by inhibiting glutamate production, leading to reduced cellular antioxidant activity. The combined supplementation of BCKA catabolites and glutathione nearly completely suppressed the cellular senescence induced by BCAT2 knockdown, indicating that both BCAA catabolites and glutamate, as products of BCAT2-mediated reactions, play important roles in suppressing cellular senescence (Figure 6F).

Whereas oxidative stress is well known to induce cellular senescence, the mechanism of BCAA catabolism leading to the inhibition of cellular senescence remains unclear. The final catabolites of BCAA metabolism are acetyl coenzyme A (acetyl-CoA) and succinyl-CoA, which enter the TCA cycle. Therefore, a reduction in BCAA catabolism during senescence might exacerbate the substrates of TCA cycle deficiency and decrease mitochondrial oxidative phosphorylation (OXPHOS). The state of mitochondrial oxidative metabolism in senescent cells has been controversial. Some reports have suggested that mitochondrial oxidative metabolism is enhanced in senescent cells ^6–8^, but these findings primarily come from studies using OIS or cancerous cell lines, in which oncogenes may alter energy metabolism. It hence remains unclear as to whether or not the upregulation of TCA cycle activity is a common characteristic of cellular senescence. In contrast, other studies have shown a reduction of OXPHOS during RS ^5,41,42^. Further research is needed to clarify the level of mitochondrial activity, including BCAA metabolism, in senescent cells.

BCAAs, particularly leucine, are also crucial for mechanistic/mammalian target of rapamycin (mTOR) signaling. Leucine activates mTORC1 (mTOR complex 1) by inhibiting Sestrin2 ^43^. mTOR signaling has been implicated in the regulation of cellular senescence ^44,45^, and BCAA-induced cellular senescence through mTOR signaling has been reported in HepG2 hepatocarcinoma cell lines ^46^. However, in our experimental conditions, the presence of excessive BCAAs did not affect the proliferation of IMR90 cells; instead, BCAA supplementation delayed the induction of senescence (Supplementary Figure S4E, F). This suggests that leucine-mediated regulation of cellular senescence via mTOR signaling may be cell type- or context-dependent, warranting further investigation.

Previous studies on *Bcat2*-KO mice have primarily focused on lipid and glucose metabolism, reflecting the epidemiological associations of BCAA with obesity and type 2 diabetes ^47^. *Bcat2* disruption in mice has been shown to reduce body weight and improve glucose tolerance, but decrease exercise tolerance ^40,48^. However, the association between *Bcat2* disruption and cellular senescence has not been investigated to date. RNA-sequencing data from the skeletal muscle of *Bcat2*-KO mice demonstrated increased expression of *Cdkn1a* (p21), *Ccl8*, *Cxcl13*, and *Ccl19* genes, indicating cellular senescence ^49^. In the present study, we demonstrated the accumulation of senescent cells in the skeletal muscle and adipose tissue of *Bcat2*-KO mice (Figure 5D–G, I–K, M, Supplemental Figure S8). The reason why cellular senescence was induced in these tissues may be linked to the metabolic rates of BCAA and the expression levels of *Bcat2*. Skeletal muscle is characterized by the highest rate of BCAA metabolism in the body ^50^. *Bcat2* expression is high in adipose tissue, particularly in white adipose tissue (Figure 6B, C).

The expression of p21 and SASP-related genes was significantly upregulated in the skeletal muscle of *Bcat2*-KO mice. In contrast, an increase in the levels of *Cdkn*s and cytokines in adipose tissue was not evident. Instead, the expression of *Mmp*s was markedly increased, suggesting that cellular senescence promotes tissue remodeling in the adipose tissue of *Bcat2*-KO mice. Our quantification using the reverse transcription real-time quantitative PCR (RT-qPCR) on bulk tissues may have underestimated these changes, owing to the increased presence of interstitial tissues, as adipose tissue atrophy was notably severe. However, recent studies have shown that the upregulation of SASP-related genes is not uniform, but demonstrates diverse patterns depending on the stimuli or cell types ^51–53^. Furthermore, increased levels of CDKNs were not consistently observed as a common signature of cellular senescence ^51^. The results of our present study may also reflect tissue-specific responses. Moreover, as cellular senescence in *Bcat2*-KO mice was induced in myocytes and adipocytes, i.e., differentiated, post-mitotic cells, the continuous expression of CDKNs might not be crucial. As most common markers of cellular senescence have been identified in dividing cells, further research is required on cellular senescence in nondividing cells *in vivo*.

Finally, BCAA metabolism, particularly its transamination, is reduced in normal aging (Figure 6). Normal aging results in decreases in *Bcat2* expression and BCKA production in mouse adipose tissue. Alterations of BCAA metabolism were also observed in humans. Taken together with the fact that intervening with BCAA intake affects lifespan in several animal experiments ^23, 26–28^, our findings indicate that the BCAA transamination reaction may be a crucial step in aging.

## Materials and methods

### Ethics approval

All animal experiments were approved by the Institutional Animal Care and Use Committee of Gunma University and Tokyo Medical and Dental University.

### Cells and reagents

RPE1-hTERT and IMR90-hTERT cells were kind gifts from Makoto Nakanishi and Makoto Hayashi, respectively. Doxycycline (Dox), gabapentin, doxorubicin, H_2_O_2_, and neocarzinostatin were purchased from Sigma-Aldrich (St. Louis, MO, USA). Antibodies against human BCAT2 (#9432) and phospho-BCKDHA (#90198) were purchased from Cell Signaling Technology (Danvers, MA, USA). Antibodies against mouse BCAT2 (A7426) and BCKDHA (A19962) were purchased from ABclonal (Woburn, MA, USA). Anti-β-actin (A1978) and anti-α-tubulin (T6074) antibodies were purchased from Sigma-Aldrich. The anti-lamin B1 antibody (12987-1-AP) was purchased from Proteintech (Rosemont, IL, USA). Isobutyryl-carnitine, isovaleryl-carnitine, and methylbutyryl-carnitine were purchased from Cayman Chemical (Ann Arbor, MI, USA).

### Cell culture

IMR90-hTERT and RPE1-hTERT cells were cultured in Dulbecco’s modified Eagle’s medium, low glucose and high glucose, respectively (Thermo Fisher Scientific, Waltham, MA, USA), containing 10% Fetal Bovine Serum (FBS), 1× GlutaMAX, 1 mM pyruvate, 1× nonessential amino acids, and 1× penicillin–streptomycin (Thermo Fisher Scientific) in a 5% O_2_ / 5% CO_2_ atmosphere. IMR90-tetTRF2ΔB and RPE1-tetTRF2ΔB cells were induced by adding 2 µg/mL Dox, and the culture medium was changed every 3 days to fresh medium containing 2 µg/mL Dox.

### Chemical induction of cellular senescence

IMR90-hTERT and RPE1-hTERT cells were treated with 200 ng/mL neocarzinostatin for 2 h, 800 µM H_2_O_2_ for 2 h, or 250 nM doxorubicin for 24 h, and incubated for 10 to 14 days. The culture medium was changed every 3 days. H_2_O_2_ treatment was applied repeatedly every 3 days. After induction, cells were subjected to the SA-β Gal staining assay to confirm cellular senescence.

### Plasmids

The lentiviral tet-inducible TRF2ΔB (amino acids 45–497) expression plasmid was constructed by cloning the complementary DNA (cDNA) of human TRF2 with an N-terminal myc-tag into the pLVX-tetOnePuro plasmid (Takara Bio, Kusatsu, Japan). The lentiviral human BCAT2 expression plasmid using a pLVXneo backbone was designed and ordered from VectorBuilder (Chicago, IL, USA). The lentiviral shRNA hairpin expression plasmids for BCAT2 and KLF15 were constructed by cloning the synthetic oligonucleotides into the pLKO.1 puro plasmid (Sigma-Aldrich). The shRNA target sequences were as follows.

Control: TCCGCAGGTATGCACGCGTG

BCAT2 #1: TGAAGTGCAATACGAAATAAA

BCAT2 #2: GTGCACCGAATCCTGTACAAA

KLF15 #1: AGTTGGGTATCTGGGTGATAG

KLF15 #2: ACATAGGTCCATCCACATAAA

### Lentiviral vector production and infection

Lenti-X 293T packaging cells (Takara Bio) were cotransfected with the desired DNA plasmids and the packaging plasmids (pCMV-VSVG, pMDLg/pRRE, and pRSV-REV). At 48, 60, and 72 h after transfection, the culture medium containing viruses was collected and used to infect cells in the presence of polybrene (4 µg/mL). At 48 h after the last infection, cells were selected with the desired drug and then maintained in the selection medium.

### SA-β Gal staining assay

The SA-β Gal-staining assay for culture cells was performed using the Senescence β-Galactosidase Kit (Cell Signaling Technology) according to the manufacturer’s instructions.

Staining of adipose tissues was performed as described previously ^54^. Briefly, fresh adipose tissue was fixed with 1× fixation buffer for 1 h. After the tissues were washed with phosphate buffer saline (PBS) three times, the sections were stained with the staining solution for 26 h at 37 °C. Then, the tissues were washed with PBS three times, and dehydrated by consecutive 30-min incubations in 70% and 100% ethanol. The samples were embedded with paraffin, and sectioned at 3-µm thickness. After paraffin removal, the sections were counterstained with eosin. Images were captured using a DMIL LED microscope (Leica Microsystems, Wetzlar, Germany).

### Immunofluorescence and image analysis

Cells grown on coverslips were fixed for 15 min in 2% paraformaldehyde at room temperature. After washing, the cells were permeabilized with 0.5% NP-40 in PBS, and then incubated for 30 min in the blocking solution (0.5% bovine serum albumin (BSA) 0.2% cold water fish gelatin in PBS). Cells were then incubated with primary antibodies in the blocking solution for 1 h at room temperature, washed three times with the blocking solution, incubated with secondary antibodies in the blocking solution for 1 h, and washed again three times in the blocking solution. Finally, slides were mounted in ProLong Gold antifade reagent with 4’, 6-diamidino-2-phenylindole (DAPI) (Thermo Fisher Scientific). Fluorescence images were acquired using an FV10i-DOC laser scanning microscope (Olympus, Tokyo, Japan). The images were processed using Photoshop (Adobe, San Jose, CA, USA). Quantitative image analyses were performed using MATLAB with Imaging Processing Toolbox software (MathWorks, Natick, MA, USA).

Frozen sections were washed with PBS and heated for 30 min in antigen retrieval buffer (20 mM Tris pH 9.0, 1 mM ethylenediaminetetraacetic acid (EDTA) 0.05% Tween 20). After cooling, the sections were permeabilized with 1% Tween 20 in PBS (PBS-T) for 5 min, washed with PBS, and blocked with 10% FBS in PBS-T for 30 min. Then, the sections were incubated with primary antibodies in the blocking buffer overnight at 4 °C. After incubation, the sections were washed with PBS three times, and incubated with secondary antibodies in Can Get Signal immunostain solution A (TOYOBO, Osaka, Japan) for 1.5 h. The sections were counterstained with DAPI, and mounted with FluorSave Reagent (Merck, Darmstadt, Germany). Images were captured using an FV10i-DOC microscope (Olympus). Analysis of nuclear lamin B1 intensity was performed using Fiji software ^55^.

### Telomere dysfunction-induced foci (TIF) assay

Immunofluorescence-fluorescence *in situ* hybridization was used to detect TIFs, as described previously ^30^, using primary antibodies against phosphorylated histone H2AX on serine 139 (γH2AX). Cells grown on coverslips were fixed for 15 min in 2% paraformaldehyde at room temperature, followed by 15 min in 100% methanol at −20 °C. After rehydration in PBS for 5 min, cells were incubated for 30 min in the Blocking Solution (1 mg/mL BSA, 3% goat serum, 0.1% Triton X-100, and 1 mM EDTA in PBS). The cells were incubated with primary antibodies in Blocking Solution for 1 h at room temperature, washed three times in PBS, incubated with secondary antibodies in Blocking Solution for 30 min, and washed again three times in PBS. Coverslips were then fixed with 2% paraformaldehyde for 15 min at room temperature, washed twice in PBS, dehydrated consecutively in 70%, 95%, and 100% ethanol for 5 min each, and allowed to dry completely. Hybridizing Solution (70% formamide, 1 mg/mL blocking reagent [Roche, Indianapolis, IN, USA], and 10 mM Tris-HCl, pH 7.2, containing the peptide nucleic acid probe FITC-OO-(CCCTAA)3 [Bio-Synthesis, Lewisville, TX, USA]) was added to each coverslip, and the cells were denatured by heating for 10 min at 80 °C on a heat block. After 2 h of incubation at room temperature in the dark, the cells were washed twice with Wash Solution (70% formamide and 10 mm Tris-HCl, pH 7.2) and twice in PBS. Slides were mounted in ProLong Gold antifade reagent with DAPI (Thermo Fisher Scientific).

### Histological analysis

Frozen sections of gastrocnemius muscle and paraffin-embedded sections of gonadal adipose tissue were cut for hematoxylin and eosin (H&E) staining. The slides were stained with H&E following a standard protocol. The cross-section area of myocytes and adipocytes were measured for more than 100 cells, using Adobe Photoshop software (Adobe, Mountain View, CA, USA).

### ROS detection

Intracellular ROS were detected using ROS Detection Cell-Based Assay Kit (DHE) (Cayman Chemical), in accordance with the manufacturer’s instructions. Briefly, cells were stained with 5 µM dihydroethidium (DHE) for 90 min at 37 °C. Oxidized DHE images were acquired using a BZ-X700 fluorescence microscope with a tetramethylrhodamine isothiocyanate (TRITC) filter (Keyence, Osaka, Japan). Intracellular ROS were quantified by the fluorescence intensity of oxidized DHE overlapping with nuclei, as identified by DAPI staining, using MATLAB with Imaging Processing Toolbox software.

### RT-qPCR

Total RNA was isolated from cultured cells using RNeasy Mini Kit (QIAGEN, Venlo, Netherlands) or from skeletal muscle and adipose tissue using ReliaPrep RNA tissue Miniprep System (Promega, Madison, WI). Reverse transcription (RT) was performed using ReverTra Ace qPCR RT Kit (TOYOBO). Real-time PCR was performed using THUNDERBIRD SYBR qPCR Mix (TOYOBO) or TB Green Premix ExTaqTM II FAST qPCR (Takara Bio) using the StepOne Plus instrument (Thermo Fisher Scientific). Relative mRNA expression was normalized to β-actin mRNA (*ACTB*) expression. The sets of PCR primers used were as follows.

human BCAT2-F: CGCTGAATGGTGTTATCCTGCC

human BCAT2-R: CAGCAACTGCTTCATGGTGATCG

human BCAT1-F: GCTCTGGTACAGCCTGTGTTGT

human BCAT1-R: TGCCAGCTTAGGACCATTCTCC

human CDKN1A-F: AGGTGGACCTGGAGACTCTCAG

human CDKN1A-R: TCCTCTTGGAGAAGATCAGCCG

human CDKN2A-F: CCAACGCACCGAATAGTTACG

human CDKN2A-R: GCGCTGCCCATCATCATG

human CXCL1-F: AGCTTGCCTCAATCCTGCATCC

human CXCL1-R: TCCTTCAGGAACAGCCACCAGT

human CXCL2-F: GGCAGAAAGCTTGTCTCAACCC

human CXCL2-R: CTCCTTCAGGAACAGCCACCAA

human IL6-F: AGACAGCCACTCACCTCTTCAG

human IL6-R: TTCTGCCAGTGCCTCTTTGCTG

human IL8-F: GAGAGTGATTGAGAGTGGACCAC

human IL8-R: CACAACCCTCTGCACCCAGTTT

human KLF15-F: GTGAGAAGCCCTTCGCCTGCA

human KLF15-R: ACAGGACACTGGTACGGCTTCA

human SREBF1-F: ACTTCTGGAGGCATCGCAAGCA

human SREBF1-R: AGGTTCCAGAGGAGGCTACAAG

human PGC1a-F: CCAAAGGATGCGCTCTCGTTCA

human PGC1a-R: CGGTGTCTGTAGTGGCTTGACT

human ACTB-F: AAGACCTGTACGCCAACACAGT

human ACTB-R: ACTCCTGCTTGCTGATCCACAT

mouse Cdkn1a-F: TCGCTGTCTTGCACTCTGGTGT

mouse Cdkn1a-R: CCAATCTGCGCTTGGAGTGATAG

mouse Cdkn2a-F: TGTTGAGGCTAGAGAGGATCTTG

mouse Cdkn2a-R: CGAATCTGCAACCGTAGTTGAGC

mouse Il1b-F: TGGACCTTCCAGGATGAGGACA

mouse Il1b-R: GTTCATCTCGGAGCCTGTAGTG

mouse Tgfb1-F: TGATACGCCTGAGTGGCTGTCT

mouse Tgfb1-R: CACAAGAGCAGTGAGCGCTGAA

mouse Tnfa-F: GGTGCCTATGTCTCAGCCTCTT

mouse Tnfa-R: GCCATAGAACTGATGAGAGGGAG

mouse Cd68-F: GGCGGTGGAATACAATGTGTCC

mouse Cd68-R: AGCAGGTCAAGGTGAACAGCTG

mouse Cxcl1-F: TCCAGAGCTTGAAGGTGTTGCC

mouse Cxcl1-R: AACCAAGGGAGCTTCAGGGTCA

mouse Mmp2-F: CAAGGATGGACTCCTGGCACAT

mouse Mmp2-R: TACTCGCCATCAGCGTTCCCAT

mouse Mmp3-F: CTCTGGAACCTGAGACATCACC

mouse Mmp3-R: AGGAGTCCTGAGAGATTTGCGC

mouse Actb-F: CATTGCTGACAGGATGCAGAAGG

mouse Actb-R: TGCTGGAAGGTGGACAGTGAGG

### Measurements of relative telomere length

Genomic DNA was extracted from IMR90-hTERT cells using Quick Gene DNA tissue kit S (Fujifilm Wako Chemicals, Osaka, Japan). Mean telomere lengths were measured by qPCR, as previously described ^56^. The total volume of the PCR mixture was 20 µL, and the final concentration of the reagents in the PCR mixture were as follows: 1× Thunderbird SYBR Green I mix (TOYOBO), 900 nM of telomere primers (telG: ACACTAAGGTTTGGGTTTGGGTTTGGGTTTGGGTTAGTGT, telC: TGTTAGGTATCCCTATCCCTATCCCTATCCCTATCCCTAACA) or 500 nM of β-globin primers (hbgu: CGGCGGCGGGCGGCGCGGGCTGGGCGGCTTCATCCACGTTCACCTTG, hbgd: GCCCGGCCCGCCGCGCCCGTCCCGCCGGAGGAGAAGTCTGCCGTT), 1× ROX reference dye, 500 mM betaine (Fujifilm Wako Chemicals), and 20 ng of genomic DNA. The thermal cycle profile was Stage 1: 95 °C for 10 min, Stage 2: 40 cycles of 20 sec at 95°C, 20 sec at 52°C, and 45 sec at 72°C with signal acquisition. Samples were normalized to β-globin, and then normalized to samples on Day 0.

### Metabolite collection from culture cells

Metabolites were extracted from cells using 80% methanol at a final concentration of 3 × 10^5^ cells/mL, by gentle shaking for 30 min at room temperature. Collected extracts were centrifuged at 10,000 *g* for 5 min, and then the supernatants were applied to Captiva ND Lipids System (Agilent Technologies, Santa Clara, CA, USA) to remove proteins and phospholipids.

### Metabolite collection from mouse blood

Female C57BL/6 mice were purchased from CLEA Japan (Tokyo, Japan). Peripheral blood samples were collected in heparinized tubes from the submandibular vein using an animal lancet (MEDIpoint, Mineola, NY, USA). Blood samples were centrifuged at 1,200 *g* for 30 min to collect plasma. Metabolites were extracted with 80% methanol by gentle shaking for 30 min at room temperature, and then centrifuged at 10,000 *g* for 5 min. Collected supernatants were then applied to Captiva ND Lipids System to remove proteins and phospholipids.

### LC-MS analysis of metabolites

LC-MS analysis was performed using an LCMS-8050 triple quadrupole mass spectrometer system (Shimadzu, Kyoto, Japan). The relative levels of metabolites in central metabolic pathways were determined using the Method Package for Primary Metabolites (Shimadzu) with a Discovery HS F5-3 column (Sigma-Aldrich), in accordance with the manufacturer’s instructions. For the measurement of BCKAs, cellular extracts were derivatized with 3-nitrophenylhydrazine in the presence of pyridine and 1-ethyl-3-(3-dimethylaminopropyl) carbodiimide for 30 min at room temperature. The reaction was stopped by adding an excess amount of formic acid. The derivatized metabolites were separated on a reversed-phase MastroC18 column (Shimadzu) using a gradient of solvents A (water with 0.1% formic acid) and B (acetonitrile with 0.1% formic acid), with a flow rate of 0.35 mL/min. The initial solvent composition was 16% B, and the following linear gradient of the solvent was applied: 16% to 25% from 0 to 6 min, 25% to 40% from 6 to 9 min, 40% to 95% from 9 to 17 min, held at 95% B for 3 min, and then returned to 16% B and maintained there for 3 min. The column was maintained at 40 °C. The transitions for selected reaction monitoring of analytes were m/z 387.15 > 145.00 [M+H]^+^ for α-KIV, and 401.00 > 220.05 [M+H]^+^ for α-KIC and α-KMV.

### Normalization of metabolome data

Metabolome data were filtered by removing metabolites with a large coefficient of variation among the triplicate experiments or the sample treatments. Each metabolite level was then divided by the total amount of all filtered metabolites for normalization.

### Metabolite set enrichment analysis (MSEA)

MSEA, which is a metabolomic version of the gene set enrichment analysis (GSEA), was performed using MetaboAnalyst 5.0 software(https://www.metaboanalyst.ca/) ^57^.

### Tabula Muris Senis dataset

The Tabula Muris dataset was downloaded from https://tabula-muris.ds.czbiohub.org/.

### Measurement of plasma BCKA levels of a human cohort

The metabolomic data of human plasma were obtained from the Japanese Multi-Omics Reference Panel (jMorp; https://jmorp.megabank.tohoku.ac.jp/) ^58^. Metabolite indices (the mean and the 95% confidence interval [CI] for each metabolite across age groups) for α-KIC (TCN000008), α-KMV (TCN000006), α-KIV (TCN000038), valine (TCN000032), isoleucine (TCN000018), and leucine (TCN000027) analyzed using nuclear magnetic resonance spectroscopy are presented.

### Generation of Bcat2 KO mice

*Bcat2* KO mice were generated by deleting exons 4 to 6 using CRISPR-Cas9 genome editing. Two guide RNAs (gRNAs) flanking exons 4 to 6, namely, Bcat2L1 targeting 5′- CTCTTAGCTCAACTCCTCAG-3′ and Bcat2R1 targeting 5′-ACATCTGTGAGGTCTTCTGG- 3′, were designed. Fertilized eggs were collected from C57BL/6J mice (Jackson Laboratory, Tsukuba, Japan), and genome editing was performed by electroporation, as previously reported ^59^. Gene deletion and wild-type alleles of the offspring were detected by PCR using the following primer sets:

Bcat2-F1: GCTTGCATGTGGTGATTTTG

Bcat2-R1: GTGCAGATCCGGACACTAGG

Bcat2-R2: CGCCTAGCAGAACGTAGCAT

Finally, deleted regions were confirmed by direct sequencing using the PCR products.

### Statistical analysis

Statistical analyses were performed by the unpaired Student *t*-test, the Wilcoxon rank sum test, one-way analysis of variance (ANOVA), or two-way ANOVA with the post hoc test using R software (https://www.R-project.org/).

### Figures and Illustrations

Figures were generated using Adobe Illustrator (https://www.adobe.com) and BioRender software (https://biorender.com).

## Acknowledgments

We thank Makoto Nakanishi at the University of Tokyo for the critical reading of the manuscript and for the RPE1-hTERT cells; Makoto Hayashi at Kyoto University for the generous gift of IMR90-hTERT cells; Ikuyo Yoshino at Tokyo Medical and Dental University for the mouse husbandry; and Kayo Suzuki and Yoko Sagara at Gunma University for the technical supports in the SA-β Gal-staining of adipose tissues and LC-MS analysis of metabolites, respectively. This work was supported by the Japan Society for the Promotion of Science (JSPS) KAKENHI grant numbers JP#23K14458 to K.I., JP#25440080, JP#17K08649, and JP#21K06852 to A.K., the Platform Project for Supporting Drug Discovery and Life Science Research (Basis for Supporting Innovative Drug Discovery and Life Science Research [BINDS]) from the Japan Agency for Medical Research and Development AMED (grant number JP21am0101120 [support no. 3226]) to I.H., and the Ministry of Education, Culture, Sports, Science and Technology MEXT Project for Promoting Public Utilization of Advanced Research Infrastructure (Program for Supporting the Introduction of the New Sharing System) grant number JPMXS0420600120.

## Author contributions

Y.A., K.I., and A.K. designed the experiments. Y.A. and K.I. performed the cell biological and biochemical experiments. A.K. developed the tetTRF2ΔB inducible system, performed lamin B1 immunofluorescence, and performed quantitative imaging and statistical analyses. H.O. performed the LC-MS measurements and analyses. A.T. and A.K. analyzed the metabolomics data. T.H., R.K., and I.H. generated the *Bcat2*-KO mice. K.I., J.H., S.A., and A.K. analyzed the *Bcat2*-KO mice. S.H. and S.S. performed mouse plasma metabolomics. S.S., T.I., and Y.A.M. provided information, reagents, and advice. A.K. conceived the study and wrote the paper, with input from all the coauthors.

---

**Supplementary Figure S1.**
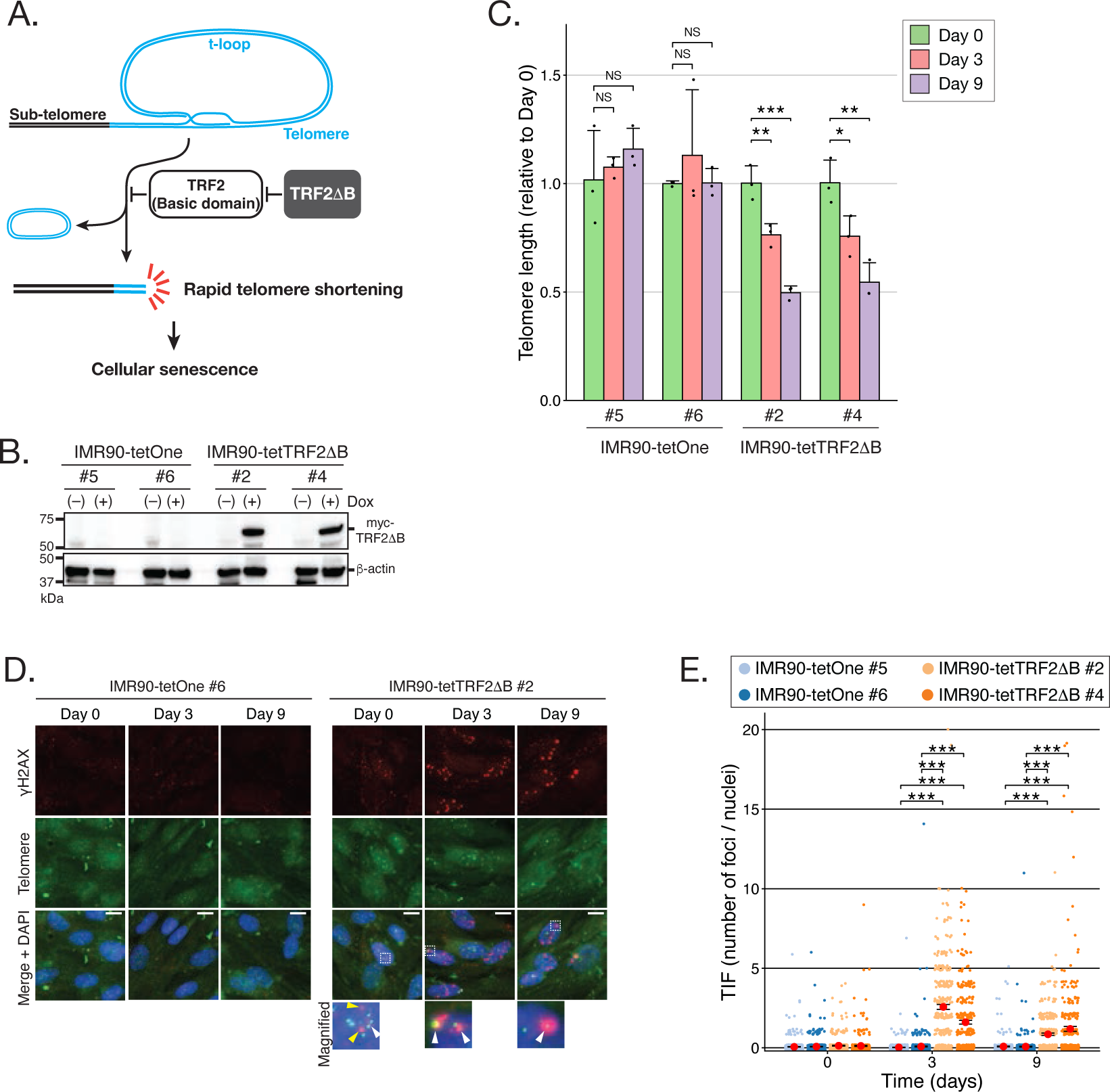
Rapid induction of telomere shortening in tetTRF2ΔB cells. **(A)** Schematic of the mechanism of telomere protection by the TRF2 basic domain. The TRF2 basic domain protects telomeres from t-loop-sized deletion. The basic domain-deletion mutant of TRF2 (TRF2ΔB) dominant-negatively inhibits TRF2, and consequently induces rapid telomere shortening and cellular senescence. **(B)** Inducible expression of the myc-tagged TRF2ΔB protein in tetTRF2ΔB cells. Two clones each of IMR90-tetTRF2ΔB cells and IMR90-tetOne control cells were incubated with or without Dox, and then subjected to immunoblotting. β-Actin was used as a loading control. **(C)** Rapid telomere shortening in Dox-induced tetTRF2ΔB cells. IMR90-tetTRF2ΔB cells or IMR90-tetOne cells were incubated with Dox for the indicated times, and then subjected to telomere length measurements. Data represent mean values of three independent experiments. Error bars indicate SD. \*\*\**p* < 0.001 and \*\**p* < 0.01, two-way ANOVA with Tukey’s test. NS, not significant. **(D, E)** Increased telomere dysfunction-induced foci (TIF) formation in Dox-induced tetTRF2ΔB cells. (D) Cells were incubated with Dox for the indicated times, and then subjected to immunofluorescence to detect γH2AX (red), and *in situ* hybridization to detect telomere DNA (green). Images at the bottom are magnified images of the boxed regions. Arrowheads indicate TIFs. Scale bars, 10 µm. (E) Quantification of TIFs in (D). Red dots indicate mean values. Error bars indicate SE. \*\*\**p* < 0.001, by two-way ANOVA with Tukey’s test.

**Supplementary Figure S2.**
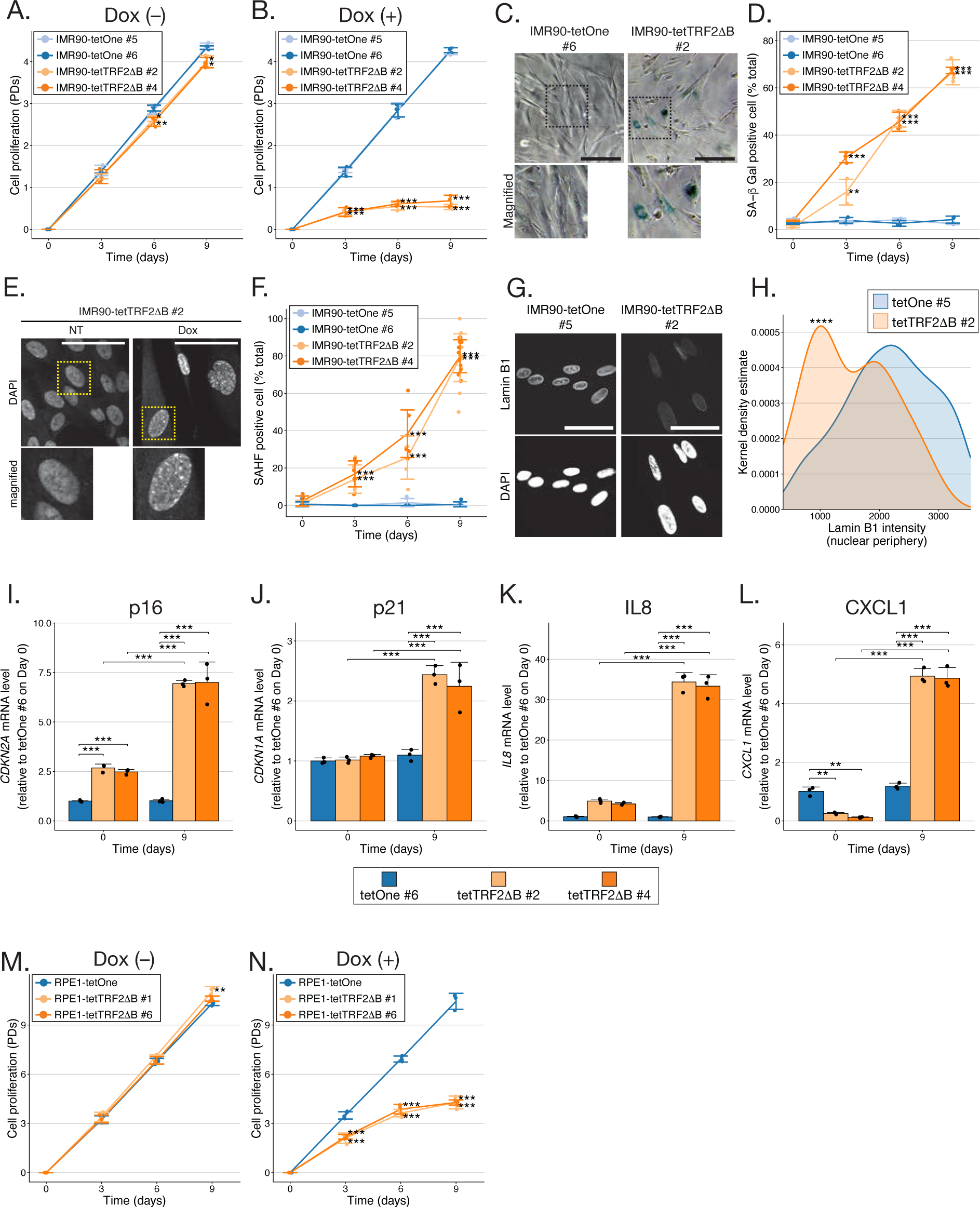
Rapid and efficient induction of cellular senescence in tetTRF2ΔB cells. **(A,B)** Rapid proliferation inhibition in Dox-treated tetTRF2ΔB cells. IMR90-tetTRF2ΔB and IMR90-tetOne cells were incubated without (A) or with Dox (B), and cumulative PDs were calculated. Data are presented as mean values from three independent experiments, and the dots represent the values from each experiment. Error bars indicate SD. \**p* < 0.05, \*\**p* < 0.01, and \*\*\**p* < 0.001; versus IMR90-tetOne #6 at each time point, by two-way ANOVA with Tukey’s test. **(C, D)** Increase of SA-β Gal activity in Dox-treated tetTRF2ΔB cells. (C) Representative images of SA-β Gal staining. IMR90-tetTRF2ΔB or IMR90-tetOne cells were incubated with Dox for 9 days, and then subjected to the SA-β Gal assay. Images at the bottom are magnified images of the boxed regions. Scale bars, 100 µm. (D) Quantification of SA-β Gal-positive cells in the experiment in (C). Data are presented as mean values from three independent experiments, and the dots represent the values from each experiment. Error bars indicate SD. \*\**p* < 0.01 and \*\*\**p* < 0.001; versus IMR90-tetOne #6 at each time point, by two-way ANOVA with Tukey’s test. **(E, F)** SAHF formation in Dox-treated tetTRF2ΔB cells. (E) Representative images of DAPI staining. IMR90-tetTRF2ΔB or IMR90-tetOne cells were incubated with Dox for the indicated times, and then subjected to DAPI staining. Images at the bottom are magnified images of the boxed regions. Scale bars, 50 µm. (F) Quantification of the SAHF-positive cells in the experiment in (E). Data are presented as mean values from three independent experiments, and the dots represent the values from each experiment. Error bars indicate SD. \*\*\**p* < 0.001; versus IMR90-tetOne #6 at each time point, by two-way ANOVA with Tukey’s test. **(G, H)** Loss of lamin B1 in Dox-treated tetTRF2ΔB cells. (G) Representative immunofluorescence images of lamin B1. IMR90-tetTRF2ΔB or IMR90-tetOne cell clones were incubated with Dox for 9 days, and subjected to immunofluorescence to detect lamin B1. Scale bars, 50 µm. (H) Quantification of lamin B1 fluorescence intensity at the nuclear periphery in the experiment in (G). \*\*\*\**p* < 0.0001, by the unpaired Student *t*-test. **(I–L)** Increased expression of *CDKN*s and SASP-related genes in Dox-treated IMR90-tetTRF2ΔB cells. IMR90-tetTRF2ΔB or IMR90-tetOne cell clones were incubated with Dox for the indicated times. Total RNAs were isolated and analyzed by qRT-PCR to detect *CDKN2A* (p16), *CDKN1A* (p21), *CXCL1*, and *IL8* mRNA. Data are normalized to β-actin, and the ratios relative to IMR90-tetOne #6 without Dox are presented. Bars represent the mean values from three independent experiments, and the dots represent the values from each experiment. Error bars indicate SD. \*\**p* < 0.01 and \*\*\**p* < 0.001, by two-way ANOVA with Tukey’s test. **(M, N)** Rapid proliferation inhibition in Dox-treated RPE1-tetTRF2ΔB cells. RPE1-tetTRF2ΔB or RPE1-tetOne cells were incubated without (A) or with Dox (B), and cumulative PDs were calculated. Data are presented as mean values from three independent experiments, and the dots represent the values from each experiment. Error bars indicate SD. \**p* < 0.05, \*\**p* < 0.01, and \*\*\**p* < 0.001; versus RPE1-tetOne #6 at each time point, by two-way ANOVA with Dunnett’s test.

**Supplementary Figure S3.**
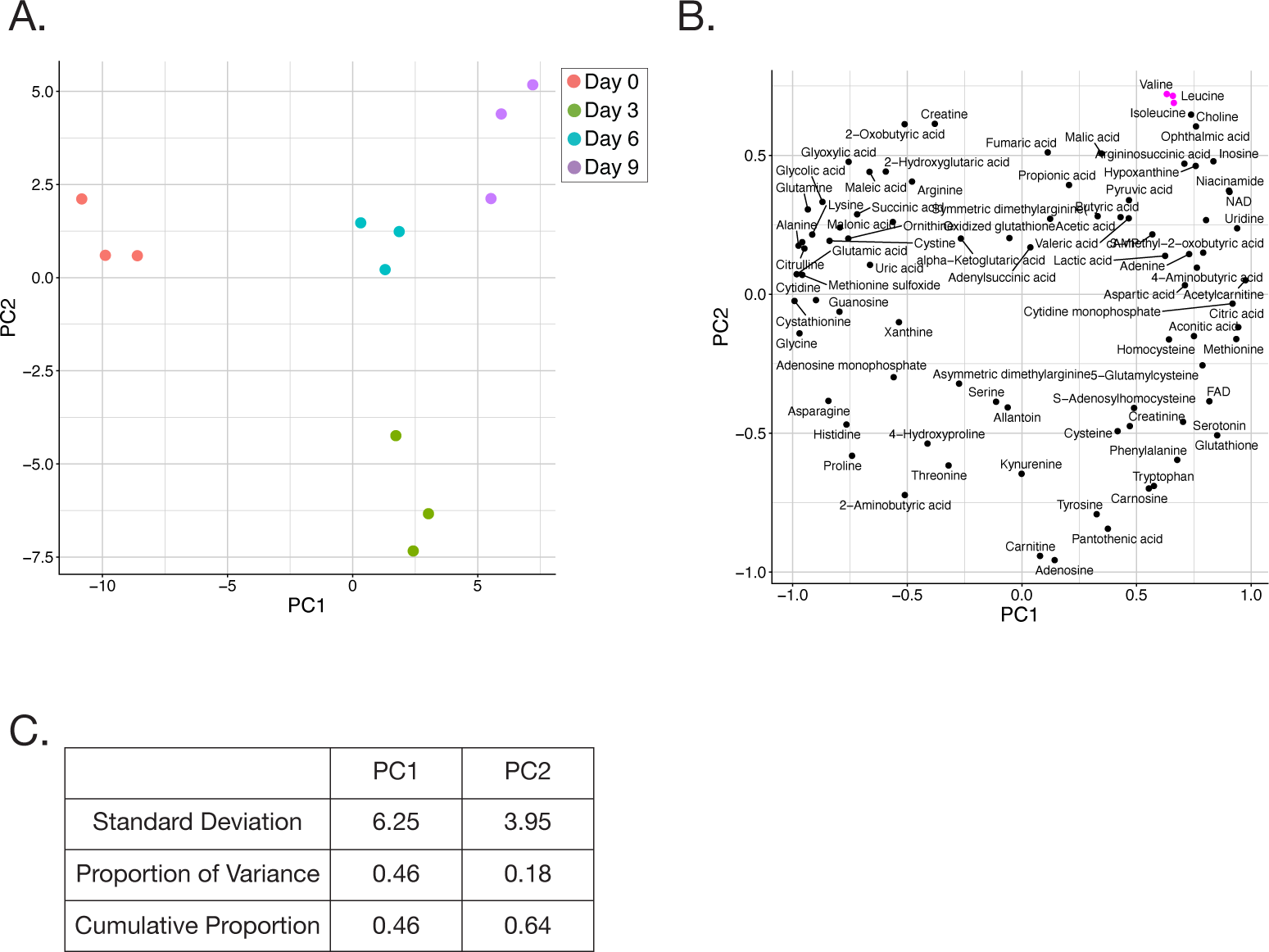
Significant shifts of the intracellular metabolome in Dox-induced tetTRF2ΔB cells. **(A–C)** Principal component analysis (PCA) of intracellular metabolites in Dox-induced IMR90-tetTRF2ΔB cells (A). Factor loading plot (B) and cumulative proportions (C) of PCA analysis. Points of the BCAAs are shown in magenta in (B). PC, principal component; FAD, flavin adenine dinucleotide; NAD, nicotinamide adenine dinucleotide.

**Supplementary Figure S4.**
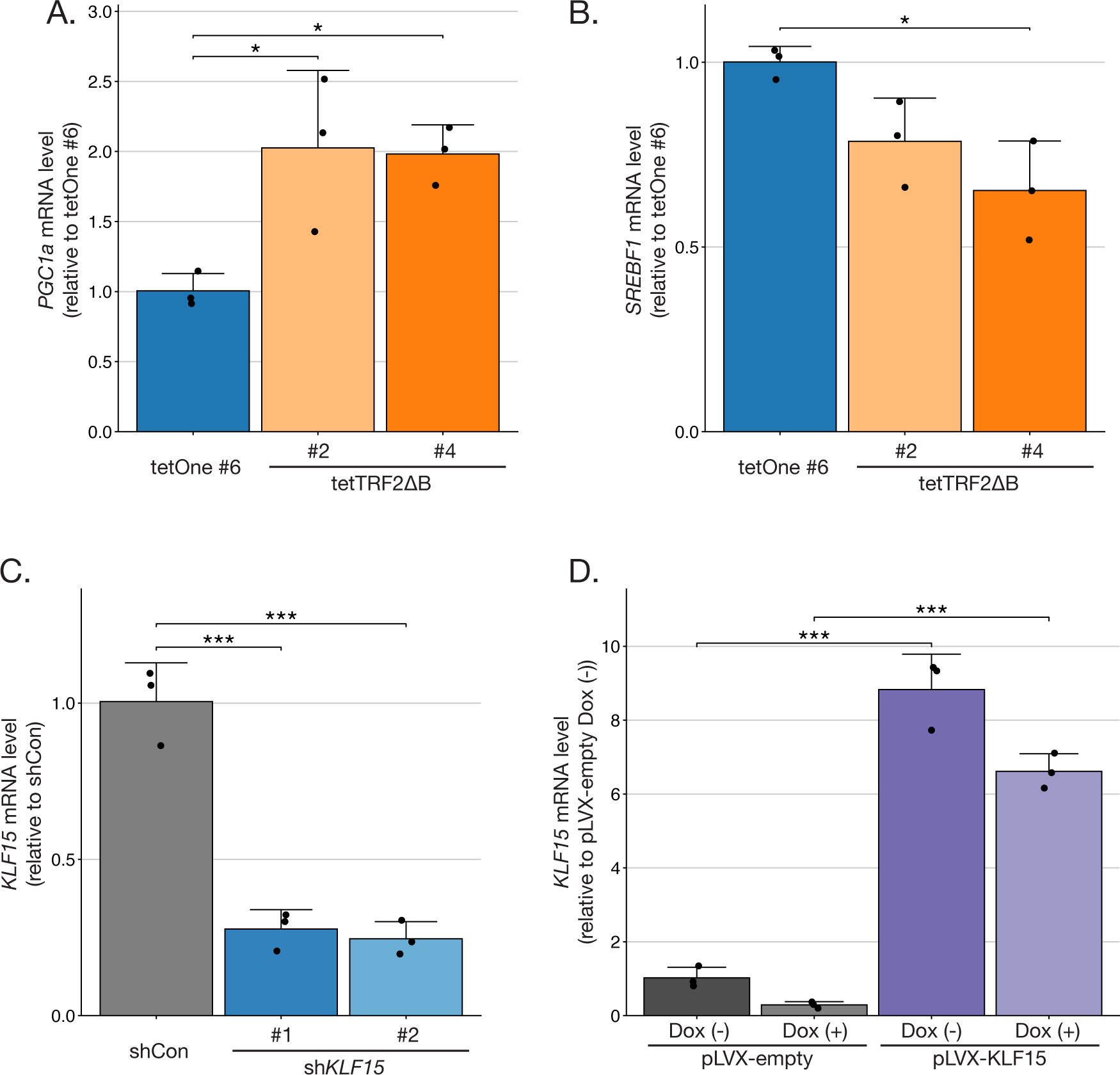
Regulation of BCAT2 suppression in senescent cells. **(A,B)** *PCG1A* and *SREBF1* expression in Dox-treated tetTRF2ΔB cells. IMR90-tetTRF2ΔB or IMR90-tetOne cells were incubated with Dox for 9 days. Total RNAs were isolated and analyzed by RT-qPCR to detect *PGC1A* (A) and *SREBF1* (B) mRNA. Data are normalized to β-actin, and the relative ratios to IMR90-tetOne #6 are presented. Bars represent the mean values from three independent experiments, and the dots represent the values from each experiment. Error bars indicate SD. \**p* < 0.05; versus IMR90-tetOne #6, by one-way ANOVA with Dunnett’s test. **(C, D)** Efficient knockdown of *KLF15* in IMR90-hTERT cells (C), and overexpression of *KLF15* in IMR90-tetTRF2ΔB cells (D). Total RNAs were isolated and analyzed by qRT-PCR to detect *KLF15* mRNA. Data are normalized to β-actin, and the relative ratios to shCon (C) or empty vector control (D) are presented. Bars represent the mean values from three independent experiments, and the dots represent the values from each experiment. Error bars indicate SD. \*\*\**p* < 0.001; versus shCon, by one-way ANOVA with Dunnett’s test (C). \*\*\**p* < 0.001; versus pLVX-empty, two-way ANOVA with Tukey’s test (D).

**Supplementary Figure S5.**
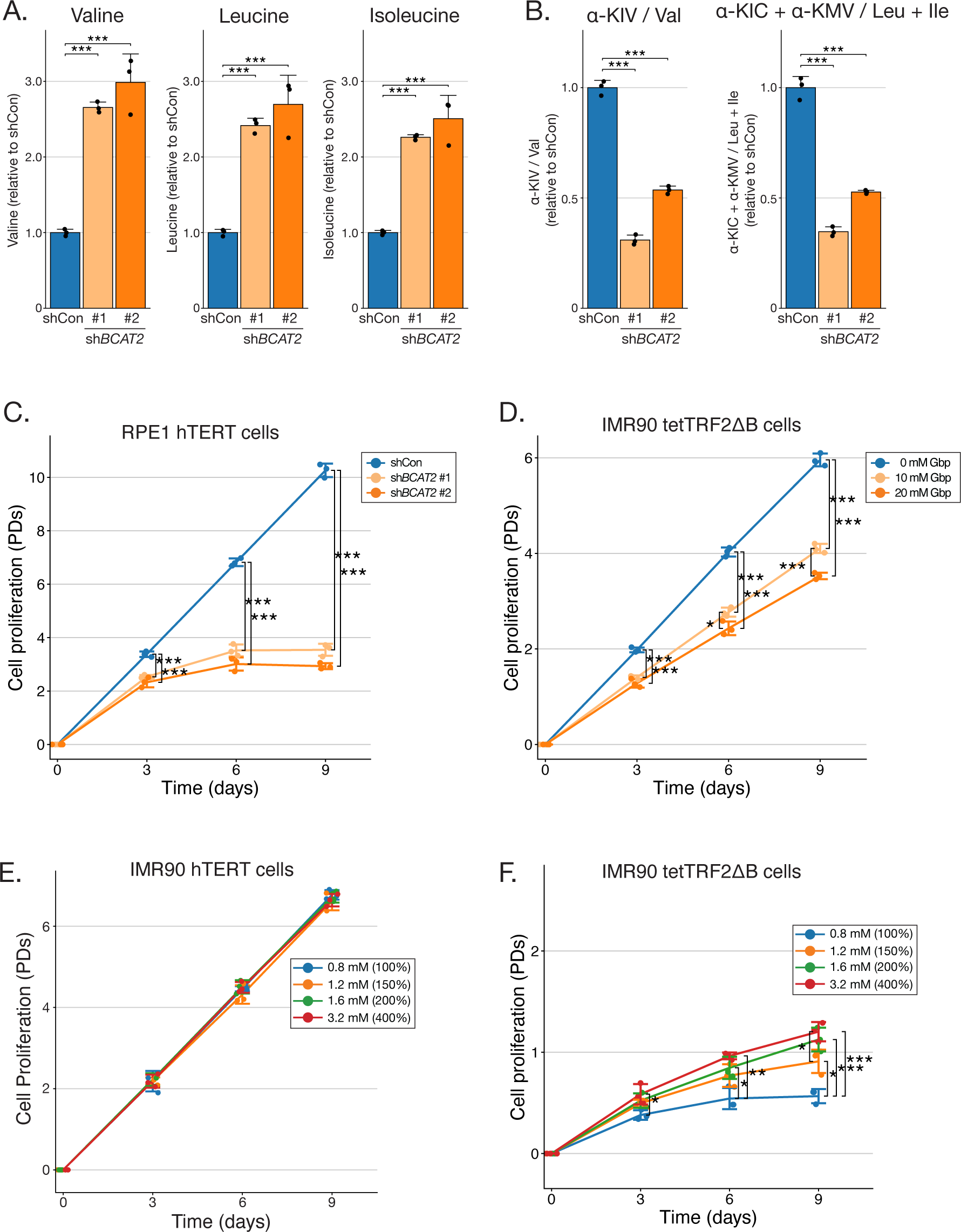
Cellular senescence induction by BCAT2 knockdown. **(A,B)** Increased BCAAs and decreased BCKA to BCAA ratios in *BCAT2*-knockdown cells. Metabolites were extracted from IMR90-hTERT cells 3 days after lentiviral shRNA introduction, and then subjected to metabolome analysis. Bars represent the mean relative ratios to shCon from three independent experiments, and the dots represent the values from each experiment. Error bars indicate SD. \*\*\**p* < 0.001; versus shCon, by one-way ANOVA with Dunnett’s test. **(C)** Rapid inhibition of proliferation in *BCAT2*-knockdown RPE1-hTERT cells. RPE1-hTERT cells lentivirally introduced with shCon or sh*BCAT2* were incubated for the indicated times, and cumulative PDs were calculated. Data are presented as mean values from three independent experiments, and the dots represent the values from each experiment. Error bars indicate SD. \*\*\**p* < 0.001, by two-way ANOVA with Dunnett’s test. **(D)** Inhibition of cell proliferation by a BCAT2 inhibitor. IMR90-hTERT cells were incubated with the BCAT2 inhibitor gabapentin (Gbp) at the indicated concentrations for the indicated times, and cumulative PDs were calculated. Data are presented as mean values from three independent experiments, and the dots represent the values from each experiment. Error bars indicate SD. \**p* < 0.05, \*\*\**p* < 0.001, by two-way ANOVA with Tukey’s test. **(E)** Cell proliferation analysis in the presence of an excess amount of BCAA. IMR90-hTERT cells were incubated with the indicated concentrations of BCAAs for the indicated times, and cumulative PDs were calculated. Data are presented as mean values from three independent experiments, and the dots represent the values from each experiment. Error bars indicate SD. **(F)** Mitigated cell growth arrest by BCAA supplementation in Dox-treated tetTRF2ΔB cells. IMR90-tetTRF2ΔB #2 cells were induced by Dox in a medium containing the indicated concentrations of BCAAs for the indicated times, and cumulative PDs were calculated. Data are presented as mean values from three independent experiments, and the dots represent the values from each experiment. Error bars indicate SD. \**p* < 0.05, \*\**p* < 0.01, \*\*\**p* < 0.001, by two-way ANOVA with Tukey’s test.

**Supplementary Figure S6.**
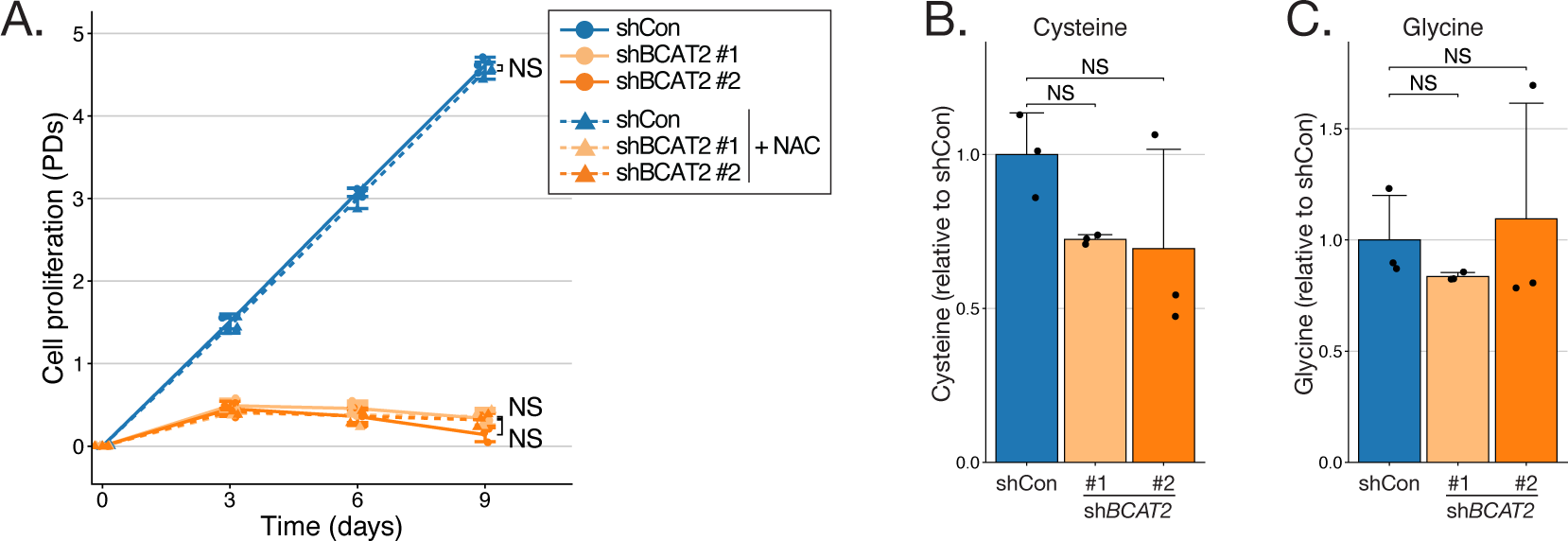
N-acetyl-L-cysteine (NAC) has no effect on cell growth arrest in *BCAT2*-knockdown cells. **(A)** Proliferation of *BCAT2*-knockdown cells incubated with NAC. IMR90-hTERT cells introduced with shCon or sh*BCAT2* were incubated with or without 5 mM NAC, and cumulative PDs were calculated. Data are presented as mean values from three independent experiments, and the dots represent the values from each experiment. Error bars indicate SD. NS, not significant, by two-way ANOVA with Tukey’s test. **(B, C)** Intracellular cysteine and glycine levels in *BCAT2*-knockdown cells. Metabolites were extracted from IMR90-hTERT cells 3 days after lentiviral shRNA introduction, and then measured by LC-MS. Metabolite levels were normalized to the total amount of all metabolites. The relative ratios to shCon are presented. Bars represent mean values from three independent experiments, and the dots represent the values from each experiment. Error bars indicate SD. NS, not significant, by one-way ANOVA with Dunnett’s test.

**Supplementary Figure S7.**
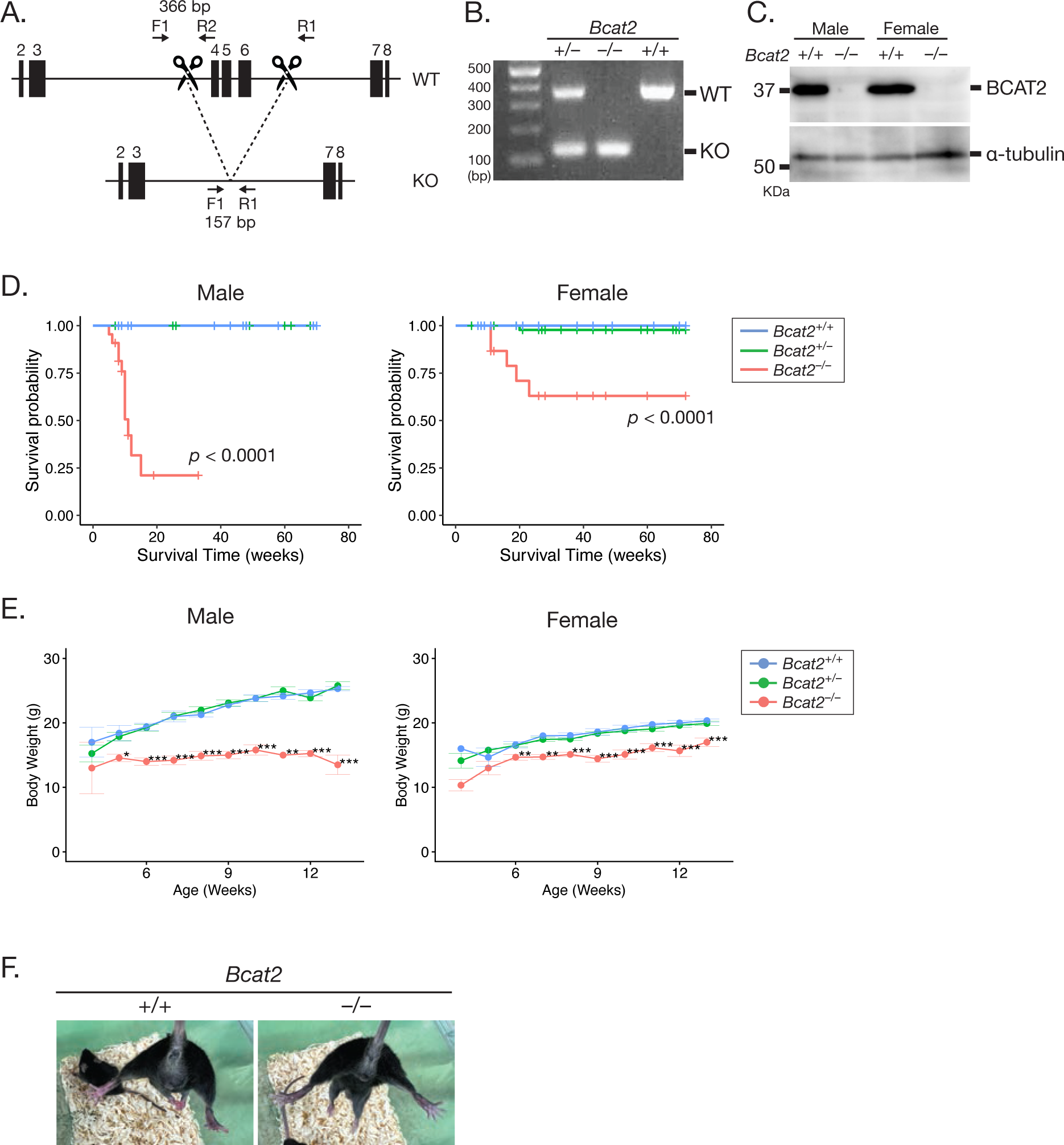
Generation of *Bcat2*-knockout mice. **(A)** Schematic showing the *Bcat2*-KO strategy using CRISPR-Cas9. Two sgRNAs were designed upstream of exon 4 and downstream of exon 6 (indicated by scissors). F1, R1, and R2, primers for PCR genotyping; bp, base pairs; WT, wild type. **(B, C)** PCR and immunoblot analysis of the indicated mice, to verify the deletion of the endogenous *Bcat2* allele (B) and protein (C), respectively. **(D)** Survival plots of *Bcat2*-KO mice. **(E)** Changes in body weight of *Bcat2*-KO mice. Error bars indicate SE. \**p* < 0.05, \*\**p* < 0.01, and \*\*\**p* < 0.001, versus Bcat2^+/+^ mice, by two-way ANOVA with Dunnett’s test. **(F)** Normal lower limb reflexes in *Bcat2*-KO mice. Representative pictures of the hindlimb clasping test of 10-week-old control and *Bcat2*-KO mice to assess signs of neurological deficits.

**Supplementary Figure S8.**
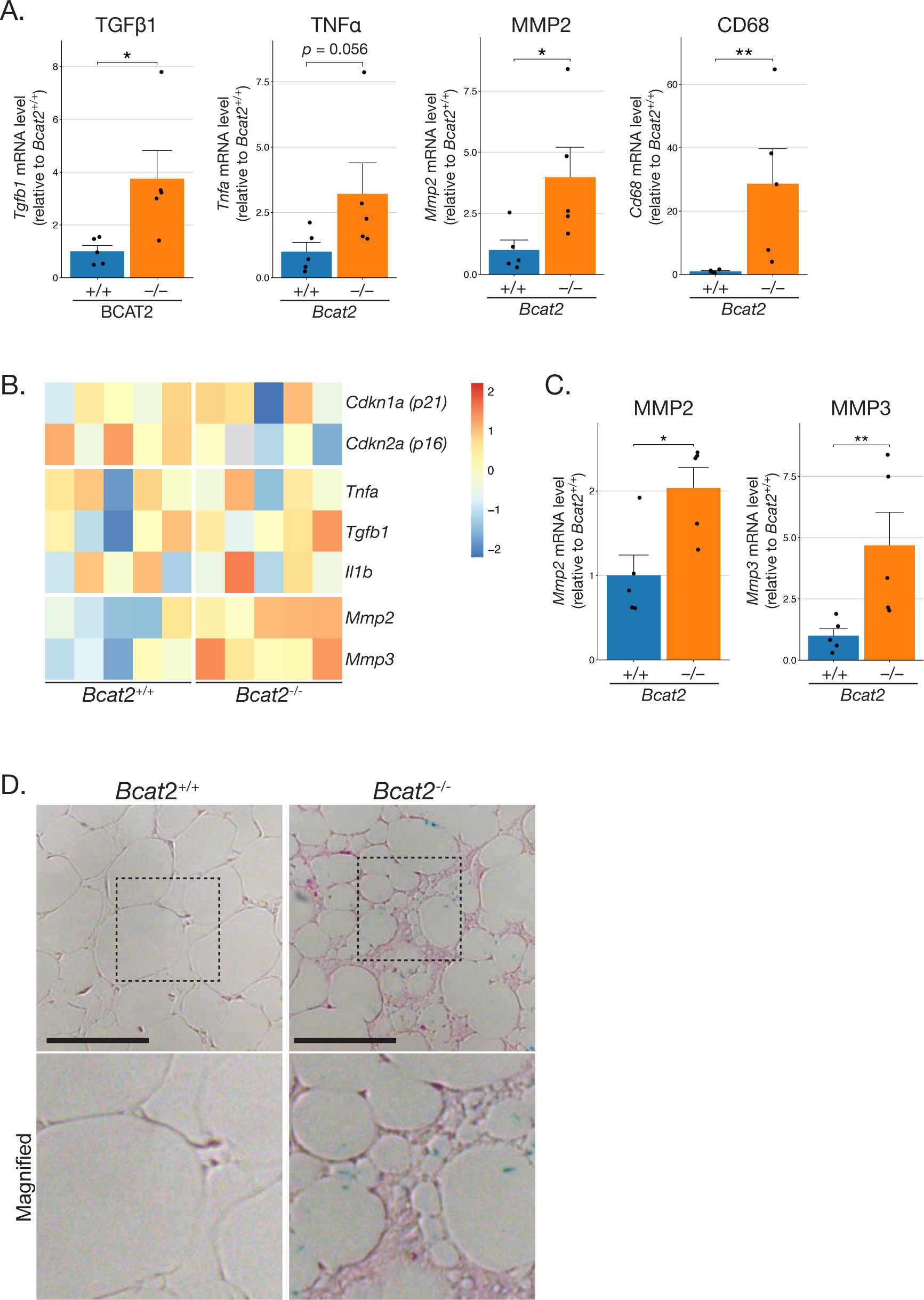
Increase in expression levels of SASP-related genes and SA-β Gal activity in *Bcat2*-knockout mice. **(A–C)** Increased expression levels of SASP-related genes in *Bcat2*-KO mice. Total RNAs were isolated from gastrocnemius muscles of 10 to 13-week-old mice (A) and gonadal adipose tissues of 10 to 11-week-old mice (B, C), shown in Fig 5F and 5G. Data are normalized to β-actin, and the relative ratios to the levels of *Bcat2*^+/+^ mice (A, C), and heatmaps of the z-scores (B) are presented. Data are presented as mean values, and the dots represent the values from each mouse (n = 5). Error bars indicate SE. \**p* < 0.05, \*\**p* < 0.01, comparing between *Bcat2*^+/+^ and *Bcat2*^−/−^ using the Wilcoxon rank sum test. **(D)** Increased SA-β Gal activity in adipocytes from *Bcat2*-KO mice. Representative histological images of SA β-Gal (blue) staining with eosin counterstaining (pink) of gonadal adipose tissues from age-matched control and *Bcat2*-KO mice (11 weeks old). The bottom images are magnified images of the boxed regions. Scale bar, 50 µm.

**Supplementary Figure S9.**
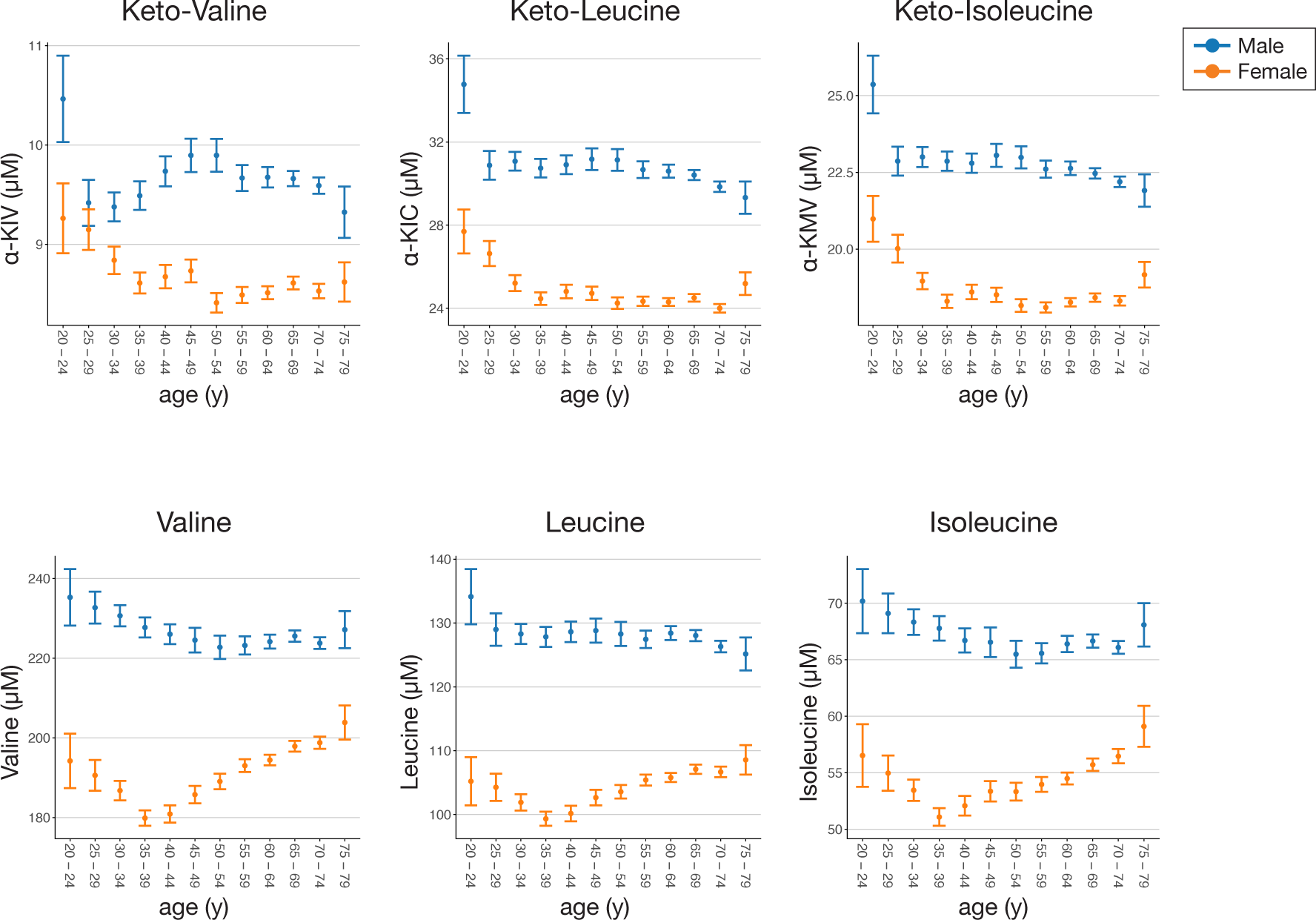
Age-associated decrease in plasma BCKA levels in a human cohort. Plasma BCKA and BCAA levels in a human cohort, grouped by age. Data represent means ± 95% CIs.

